# Pangenome analysis reveals the evolutionary dynamics of repeat-based holocentromeres

**DOI:** 10.64898/2026.01.17.700053

**Authors:** Piotr Włodzimierz, Estela Perez-Roman, Amanda Souza Câmara, Laura A. Robledillo, Gokilavani Thangavel, Meng Zhang, Jacob González Isa, Letícia Maria Parteka, Marco Castellani, Bruno Huettel, André L. L. Vanzela, Ian R. Henderson, Alexandros Bousios, André Marques

## Abstract

Centromeres are essential for chromosome segregation, yet their organisation and evolution remain poorly understood in holocentric species, where kinetochore activity is distributed along entire chromosomes^1,2^. While monocentric centromeres are often structured by megabase-sized satellite arrays^3–5^, the role of repetitive DNA in holocentric systems remains enigmatic. Here, we analyse the dynamics of centromeric *Tyba* satellite DNA repeats and transposable elements across a chromosome-scale pangenome comprising 56 long-read haplotype assemblies from 20 *Rhynchospora* species^6,7^, a plant genus with repeat-based holocentromeres^8,9^. We identify over 4.6 million monomers of the *Tyba* satellite repeat, arranged into 43,400 discrete arrays that span all chromosomes. CENH3 ChIP-seq reveals that, unexpectedly, the same *Tyba* satellite defines holocentromere across the entire genus, demonstrating deep conservation of centromeric DNA over over 40 million years despite extensive karyotype evolution and centromere array turnover. We show that *Tyba* arrays function as modular centromeric units whose number and spacing, but not size, scale with chromosome length. *Tyba* sequence diversity recapitulates species phylogeny, while higher-order repeat formation and antagonism with transposable elements shape array turnover. A novel synteny-aware algorithm reveals rapid gain, loss, and rearrangement of arrays across homologous chromosomes. Using cytogenetics and polymer simulations, we demonstrate that inter-array spacing governs chromatin loop length and chromatid thickness, linking repeat-based holocentromere organisation directly to chromosome mechanics. Our findings uncover a scalable, modular logic for holocentromere function and establish a framework for understanding the plasticity of repeat-based centromere evolution and genome architecture in eukaryotes.

## Introduction

In most eukaryotes, centromere function is confined to a single, localised region of the chromosome, a configuration known as monocentrism. However, in multiple animal and plant lineages, evolution has converged on an alternative solution: holocentricity, where kinetochore activity is dispersed almost along the entire length of the chromosome^10–12^. This architecture presents a striking departure from the canonical model, raising fundamental questions about how centromeres are structured, inherited, and evolve when uncoupled from a fixed locus.

In monocentric chromosomes, centromeres are typically organised around megabase-scale arrays of satellite repeats or transposable elements (TEs)^3–5^, bound by the centromere-specific histone CENH3 (also known as CENP-A). Recent studies have begun to elucidate the evolutionary dynamics of repeat-based centromeres in monocentric species^13,14^. Despite their ubiquity, these repeats are not strictly required for centromere function, cases of repeat-less centromeres exist, suggesting that centromeric identity is primarily epigenetically defined^15^. Holocentromeres are similarly diverse. In organisms such as *Caenorhabditis elegans* and *Bombyx mori*, holocentromeres lack canonical repeats and rely on chromatin-based cues to define kinetochore regions^16–18^. In contrast, some holocentric plants deploy satellite DNA to organise dispersed centromeric activity^8,9,19,20^, indicating that repeat-based centromere architecture can be compatible with holocentric chromosome structure. However, the mechanisms governing the assembly, stability, and evolution of repeat-based holocentromeres remain largely unknown, leaving a critical gap in our understanding of centromere diversity.

The beak-sedge genus *Rhynchospora* (Cyperaceae) has emerged as a model for repeat-based holocentromeres, as *Tyba* repeats were the first centromere-specific satellite DNA identified in a holocentric organism^8^. *Rhynchospora* chromosomes carry hundreds of short *Tyba* satellite arrays (∼20 kbp), regularly spaced and interspersed across euchromatin, which co-localise with CENH3^8,9,21^ (**Fig. 1a**). This dispersed, modular organisation differs markedly from the large, clustered centromeric arrays typical of monocentric genomes^3,9^. This distribution may be partially driven by their association with TCR1 Helitron transposons, which facilitate repeat mobility and dispersal^8,9^. Moreover, *Tyba* repeats seem to be conserved across the *Rhynchospora* genus and their abundance also inversely correlates with LTR-retrotransposon content^22,23^, suggesting a competitive relationship within the genome’s repeat landscape. Despite remarkable karyotype plasticity with frequent chromosome rearrangements and variation in chromosome number and size^24,25^, genome-wide gene synteny is largely conserved across *Rhynchospora*, hinting at unique rules governing centromere–chromosome co-evolution^6^.

**Fig. 1:**
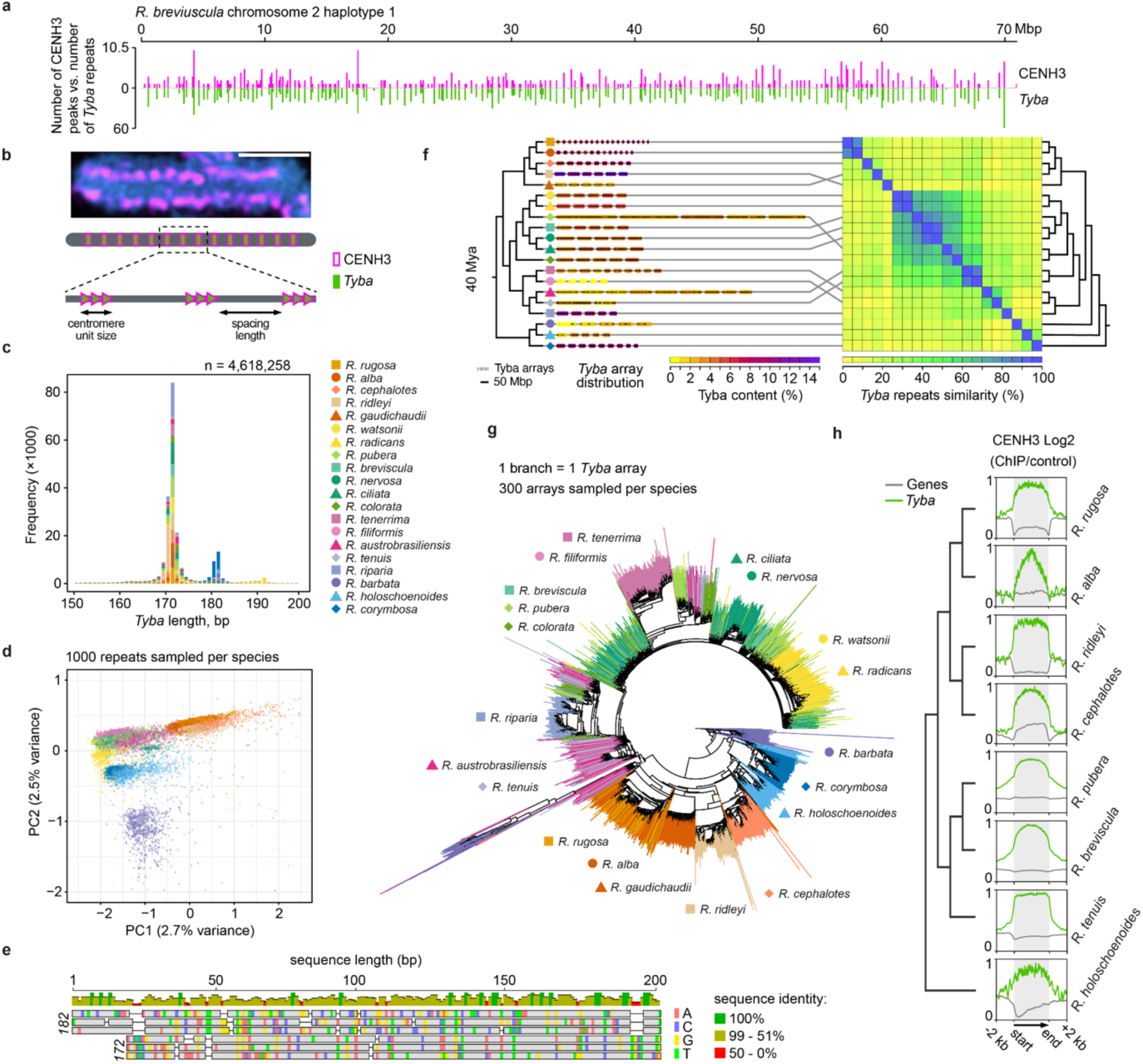
Holocentromeric organisation and sequence diversity of *Tyba* satellite DNA in *Rhynchospora*. (**a**) Chromosomal distribution of CENH3 peaks and *Tyba* repeats along chromosome 2 of *R. breviuscula* (haplotype 1), showing their specific colocalization along the entire chromosome. (**b**) A detailed view of a *Rhynchospora tenuis* mitotic chromosome stained with CENH3 and a schematic showing centromere unit size and spacing length distribution along a typical *Rhynchospora* chromosome. Scale bar corresponds to 2 µm. (**c**) Histogram of size distribution of *Tyba* repeat monomers, with a sharp peak at 172 bp and a smaller second one at 182 bp (n = 4,618,258). (**d**) Principal component analysis (PCA) of *Tyba* arrays, showing limited but structured variance across species (PC1: 2.7%, PC2: 2.5%). (**e**) Sequence alignment comparison between the 172 bp and the 182 bp *Tyba* variants. (**f**) Gene-based OrthoMCL phylogeny for the analysed *Rhynchospora* species (left) contrasted with the *Tyba* repeat similarity across species pairs (right), showing distinct clustering by phylogenetic relationship. Chromosome number and size are shown on the horizontal axis and are coloured based on their *Tyba* content. (**g**) Phylogenetic relationships among *Rhynchospora* species included in this study, with a schematic representation of *Tyba* array sampling (up to 300 arrays per species). (**h**) Metaplot of CENH3 ChIP–seq enrichment over *Tyba* arrays (green) compared to genes (grey) in eight representative *Rhynchospora* species. ChIP-seq signals are shown as log2 (normalised RPKM ChIP/input).

A key theoretical framework for explaining karyotype evolution in holocentric lineages is the holokinetic drive model^26^. It proposes that chromosome size influences total kinetochore activity during asymmetric meiosis, creating transmission biases that favour either smaller or larger homologues. These biases can drive cycles of chromosomal fission accompanied by repeat loss, or chromosomal fusion accompanied by repeat proliferation. Analogous to centromere drive in monocentric species^27,28^, holokinetic drive provides a mechanistic explanation for the divergent evolution of chromosome size and number in holocentrics.

Here, we use a chromosome-scale pangenome of 20 *Rhynchospora* species to uncover the structural logic and evolutionary dynamics of satellite-based holocentromeres. We analyse 56 long-read genome assemblies^6,7^, annotate TEs and satellite repeats, and map over 4.6 million *Tyba* monomers into 43,400 discrete centromeric arrays. CENH3 ChIP-seq confirms *Tyba*’s conserved role in centromere specification. *Tyba* arrays act as modular centromeric units: while their monomer length and array size remain stable, the number and spacing of arrays scale with chromosome length. Despite rapid sequence turnover, *Tyba* diversity reflects species phylogeny and is shaped by antagonistic interactions with TEs, consistent with a model of holokinetic drive. *Tyba* higher-order repeat formation resembles patterns seen in monocentric centromeres, suggesting convergent mechanisms of array homogenisation. We apply a new bidirectional array-matching algorithm that reveals frequent gains, losses, splitting and merging events at syntenic loci. Polymer modelling and cytogenetics demonstrate that inter-array spacing dictates chromatin loop length and chromatid thickness, directly linking repeat-based holocentromere organisation to chromosome mechanics. Together, our findings reveal a scalable, repeat-based framework for holocentromere organisation in *Rhynchospora*. By demonstrating how satellite repeats can form stable yet flexible modular units, this work provides mechanistic insights into the plasticity of centromere function, genome architecture, and the evolution of eukaryotic chromosome structure.

## Results

### Organisation and diversity of holocentromeric repeats in

#### Rhynchospora

In *Rhynchospora*, holocentromeres span entire chromosomes as regularly spaced *Tyba* repeat arrays co-localised with CENH3 by ChIP-seq^8,9,21^. These arrays show consistent spacing and form the basis of a modular centromere architecture (**Fig. 1a–b**). To investigate the distribution and evolutionary patterns of *Tyba* satellite DNA arrays in *Rhynchospora*, we developed a custom pangenome framework to annotate TEs, genes, and satellite repeats using EDTA, DANTE, Helixer, and TRASH2 (**Supplementary Fig. 1**), respectively, across 56 high-quality *Rhynchospora* haplotypes from 20 different species (**Supplementary Table 1**)^6,7^.

Remarkably, *Tyba* repeats were identified in all analysed species, comprising over 40 million years of evolution (**Fig. 1c–f**). In contrast, other satellite repeats were only found in closely related species (**Supplementary Fig. 2a**). Histogram analysis of *Tyba* monomer sizes across over 4.6 million *Tyba* monomers recovered a dominant 172 bp length and a minor but phylogenetically restricted 182 bp variant, the latter being specifically found in the earliest branching clade composed of *R. barbata*, *R. holoschoenoides*, and *R. corymbosa* (**Fig. 1c**, **Supplementary Datasets 1 and 2**). Principal component analysis of 20,000 representative monomers showed private clustering of sampled repeats from individual species (**Fig. 1d**). Apart from lower similarity levels with the dominant 172 bp variant, a conserved 16/6 bp insertion/deletion can be identified at the beginning of the 182 bp monomer variant, suggesting a single event behind the divergence and lineage-specific expansion (**Fig. 1e**).

Next, we identified over 43,400 *Tyba* arrays that are widely distributed across all chromosomes in all *Rhynchospora* species analysed, except in *R. filiformis*, which presented a genome largely with almost no *Tyba* repeats (**Fig. 1f and Supplementary Fig. 2**). Despite extensive karyotype evolution in the genus *Rhynchospora* due to frequent chromosome fissions and fusions, a high colinearity of syntenic blocks is largely found across the genus phylogeny^6^. Remarkably, gene-based phylogeny and phylogenetic relationships calculated using *Tyba* monomer sequences, using Levenshtein distance-based similarity matrix, revealed a high concordance (**Fig. 1f**). Similarly, when repeats from centromeric arrays are sampled and pairwise compared through neighbour-joining, they are arranged on the distance tree according to their expected phylogenetic relationship (**Fig. 1g**)^22^. Interestingly, several species groups clustered together, for example, *R. austrobrasiliensis* with *R. tenuis* and *R. breviscula* with *R. pubera* and *R. colorata*, indicating close relationships which might allow for a fine evolutionary analysis. These patterns demonstrate that *Tyba* sequence evolution retains a phylogenetic signal, despite pervasive array turnover.

Notably, *R. filiformis* represents an exceptional case, with only a single *Tyba* array containing 254 repeats and few other short *Tyba* stretches found elsewhere in the genome (**Fig. 1f and Supplementary Fig. 2b**). Despite this apparent loss of *Tyba*-driven holocentromeres, the residual array retains full sequence similarity to its close relative *R. tenerrima* in both array-based and repeat-based phylogenies, suggesting a very recent *Tyba* purging event in *R. filiformis* (**Fig. 1g** and **Supplementary Fig. 2b–c**). Intriguingly, no alternative putative holocentromeric repeat could be identified in the *R. filiformis* genome (**Supplementary Fig. 2**).

To investigate why *R. filiformis* no longer retains canonical *Tyba* arrays, we first examined the CENH3 sequence and a broader set of kinetochore proteins across all species. All core kinetochore components were conserved, including in *R. filiformis*. We identified a single *KNL2* copy^29^ and, as an exceptional case, lineage-specific duplications of *CENP-C*,^30^ which is uncommon in plants but should not disrupt overall kinetochore architecture^31^ (**Supplementary Fig. 3**). Notably, an entropy-based analysis of multiple sequence alignments suggested that CENP-C, CENH3 and KNL2 might evolve significantly faster than the mean across the kinetochore set (**Extended Data Fig. 1**), consistent with long-standing observations that centromere proteins are among the most rapidly evolving components of the chromosome segregation machinery^32^. However, despite this accelerated divergence, no species, including *R. filiformis*, displayed loss or extreme modification of these proteins. These findings indicate that the unusual absence of *Tyba* repeats in *R. filiformis* cannot be attributed to degeneration of core centromere proteins, and instead suggests a recent loss of the *Tyba* satellite itself.

We next performed CENH3-ChIP-seq across eight representative *Rhynchospora* species spanning all major clades of the genus, to validate the specific association of *Tyba* with CENH3. In every case, *Tyba* sequences co-localised with CENH3 binding peaks (**Fig. 1h**), confirming their conserved centromeric function over approximately 40 million years of evolution^22^. This consistent association allows *Tyba* to serve as a reliable proxy for studying repeat-based holocentromere evolution across the *Rhynchospora* pangenome, and the striking variation in *Tyba* abundance among closely related species highlights its utility as a marker for fine-scale centromere evolutionary change.

### A hierarchical higher-order repeat formation

To investigate the diversity and homogenisation of *Tyba* satellite repeats, we quantified Higher Order Repeat (HOR) scores across the 20 *Rhynchospora* species. *Tyba* repeats were binned across arrays, chromosomes, genomes, and species to evaluate patterns of sequence similarity at multiple hierarchical levels. When averaged across individual arrays, patterns similar to monocentric centromeres emerged^3^, where higher levels of HOR accumulation were found towards the centres of arrays (**Fig. 2a**). This indicates that mechanisms involved in centromeric array expansion and contraction might be parallel in monocentric and holocentric species with satellite-based centromeres, where individual arrays behave as single centromeric units. Additionally, species with the greatest HOR accumulation also had the highest genome-wide *Tyba* content (p < 0.00006, **Fig. 2b**), whereas HOR levels were negatively correlated with TE abundance (p < 0.00192) and chromosome size (p < 0.04763, **Fig. 2c–d**). These relationships are consistent with genomes bearing shorter chromosomes being enriched for *Tyba* and HORs, implicating *Tyba* homogenisation and restrained TE expansion in the maintenance and evolution of chromosome size. Across all species, *Tyba* repeat similarity is highest within individual arrays, indicating local homogenisation processes acting at the scale of single centromeric units. At the chromosomal level, similarity decreases, reflecting diversification between dispersed arrays on the same chromosome. Notably, chromosome- and genome-wide similarity are indistinguishable, indicating that each *Tyba* array evolves largely independently, with reduced homogenisation occurring beyond the array itself (**Fig. 2e**). Although this hierarchical pattern was conserved across *Rhynchospora*, the degree of intra-array homogenisation varied among species, as exemplified by the contrasting profiles of *R. rugosa* and *R. tenuis* (**Fig. 2f–g; Extended Data Fig. 2 and 3**). A small fraction of highly similar repeats was nevertheless shared across chromosomes and even between species, suggesting rare exchange events or the persistence of ancestral variants. Statistical comparisons confirmed significant differences between similarity distributions at array, chromosome, and genome levels (p < 0.001), supporting a model in which homogenisation operates locally while sequence turnover dominates at broader genomic scales. Together, these results show that *Tyba* satellites act as modular units, highly homogenised within arrays but rapidly diverging across chromosomes, reflecting a balance between local cohesion and global diversification in holocentric centromeres.

**Fig. 2:**
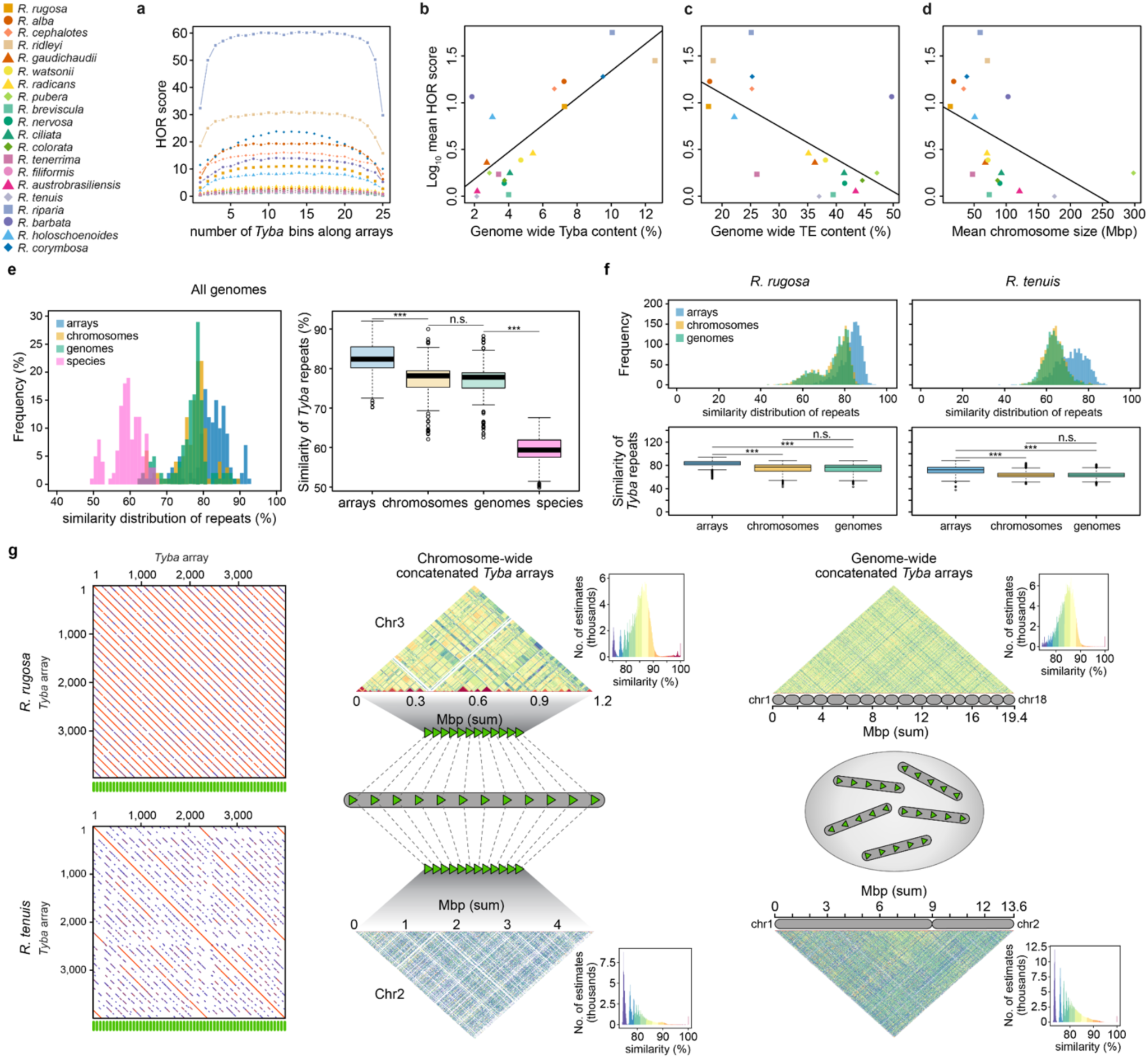
Hierarchical homogenisation of *Tyba* repeats across genomic scales in *Rhynchospora*. (**a**) Higher-order repeat (HOR) scores plotted along *Tyba* arrays, showing elevated HOR values towards array centres across species. (**b–d**) Relationship between mean HOR levels and genome-wide *Tyba* (**b**) and TE (**c**) content and mean chromosome size (**d**). (**e**) Similarity distributions of *Tyba* monomers calculated at four hierarchical levels: individual arrays, chromosomes, genomes, and across all the species analysed. Within-array repeats exhibit the highest similarity, whereas similarity decreases progressively at chromosome and genome levels, indicating strong local homogenisation and global diversification. Clearly, similarity among species is the lowest as expected. (**f**) Similarity distributions of *Tyba* monomers calculated at three hierarchical levels: individual arrays, chromosome-wide, and genome-wide in two contrasting species, *R. rugosa* (the highest chromosome number) and *R. tenuis* (the lowest chromosome number). Statistical significance was assessed using Welch’s t-tests (\*\*\**p < 0.001*; n.s., not significant). (**g**) Array (dot plot, left), Chromosome- (heatmap, middle) and genome-wide (heatmap, right) similarity plots of concatenated *Tyba* arrays in *R. rugosa* and *R. tenuis*, highlighting heterogeneous levels of local homogenization along and between chromosomes.

**Fig. 3:**
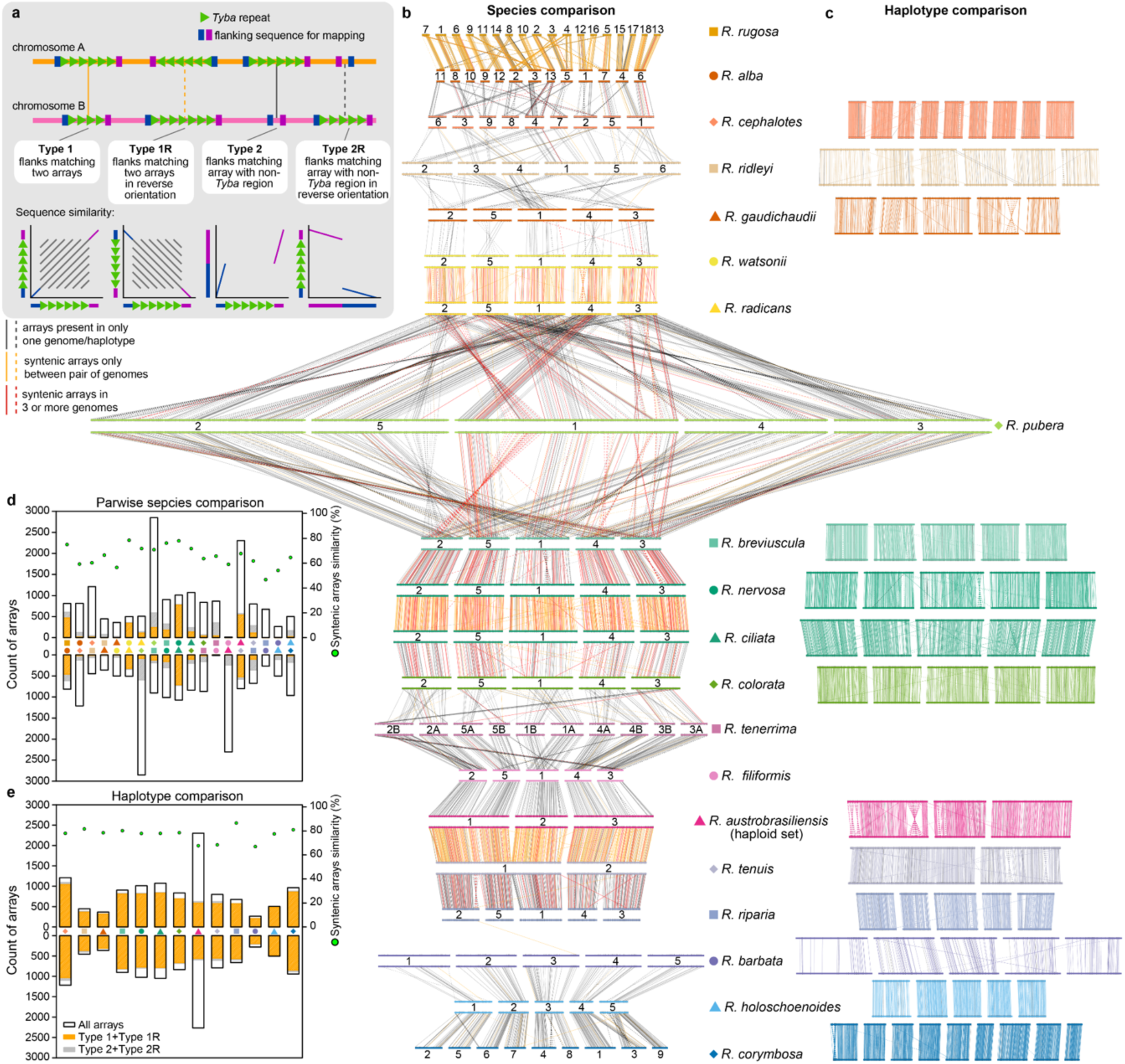
Synteny-guided tracking reveals extensive turnover and positional dynamics of *Tyba* arrays across the *Rhynchospora* pangenome. (**a**) Overview of the bidirectional array-matching strategy combining *Tyba* repeat similarity with upstream and downstream flanking context. This approach in (**b**) and (**c**) identifies orthologous/homologous arrays, detects matches to non-*Tyba* regions, and distinguishes same-strand (full line) from reversed configurations (dotted line). **Type 1** and **Type 1R** matches denote orthologous *Tyba* arrays sharing flanking sequences in the same or reverse orientation, respectively, whereas **Type 2** and **Type 2R** matches identify *Tyba* arrays aligned to syntenic non-*Tyba* regions in the same or reverse orientation, indicating array gain/loss without disruption of local genomic context. (**b**) Pairwise species comparisons of *Tyba* arrays mapped onto chromosome-scale synteny blocks across 20 *Rhynchospora* species. Coloured horizontal bars represent chromosomes, numbered according to synteny groups, and connecting lines indicate shared arrays or flanking regions, revealing conserved positioning alongside lineage-specific gains, losses and rearrangements. Syntenic arrays between pair of genomes only are highlighted as orange lines, while arrays which show synteny to three or more genomes are highlighted as red lines. Black lines correspond to an array found within a genome only. Full line and dotted line correspond to same and reverse strand, respectively. Species relationships are based on the established pangenome synteny and phylogeny from Zhang et al. ^6^. (**c**) Haplotype-level comparisons illustrating the stability of array positions within individuals, with variation largely restricted to differences in array loss or gain. Syntenic arrays between haplotypes only are highlighted as coloured lines. Black lines correspond to haplotype-specific arrays. Full line and dotted line correspond to same and reverse strand, respectively. (**d**) Quantification of shared and genome-specific syntenic loci with *Tyba* arrays across pairwise species comparisons. Please note that the total number of arrays and the proportion of syntenic arrays varies substantially, indicating rapid centromeric turnover. (**e**) Equivalent analysis for haplotype comparisons, demonstrating high positional conservation of arrays within species but some copy-number variation. Source data available in **Supplementary Dataset 3 and 4**.

### Synteny-guided tracking reveals rapid *Tyba* array turnover

To characterise the evolution of holocentromeric *Tyba* arrays across the *Rhynchospora* pangenome, we developed a bidirectional array-matching approach that integrates *Tyba* sequence similarity with flanking genomic synteny (**Fig. 3a; see Methods**). This framework identifies orthologous arrays within conserved synteny stretches (5 kbp up- and downstream arrays) and enables reconstruction of array gains, losses, positional shifts, and splitting or merging events. In this framework, **Type 1** and **Type 1R** matches correspond to orthologous *Tyba* arrays sharing conserved upstream and downstream flanking sequences in the same or reverse orientation, respectively, whereas **Type 2** and **Type 2R** matches identify *Tyba* arrays aligned to syntenic non-*Tyba* regions in the same or reverse orientation (**Fig. 3a**), indicating array loss without disruption of local genomic context. Using this approach, we tracked synteny relationships for 11,688 unique *Tyba* arrays across 20 species (**Fig. 3b, Supplementary Dataset 3**).

Despite extensive chromosome-scale synteny across the genus^6^, the centromeric landscape defined by *Tyba* arrays is highly dynamic. Closely related species often retain collinear patterns of arrays yet differ markedly in the number, size, and presence of individual units. For example, *R. alba* and *R. rugosa* maintain strong array orthology even after extensive karyotype restructuring by chromosome fissions, whereas closely related species with similar karyotypes, i.e., *R. watsonii* versus *R. radicans* or *R. nervosa* versus *R. ciliata* show pronounced synteny of *Tyba* arrays, but also cases of array losses in otherwise conserved genomic synteny (**Fig. 3b; Supplementary Figure 4**). These observations reveal that centromeric units can be gained or removed without perturbing the underlying chromosome synteny.

At the haplotype level, *Tyba* arrays are considerably more stable (**Fig. 3c, Supplementary Fig. 5; Supplementary Dataset 4**). Across multiple species, array positions are largely conserved within individuals, while copy number and spacing vary. In *R. tenuis*, comparisons among multiple accessions revealed near-complete conservation of array positions along chromosomes 1 and 2, despite minor rearrangements and copy-number differences (**Fig. 3c, Extended Data Fig. 4a**). In contrast, *R. austrobrasilensis* and *R. gaudichaudii* display extensive internal rearrangements between phased haplotypes, including inversions and reshuffling of array order, yet most arrays remain traceable across haplotypes (**Fig. 3c, Extended Data Fig. 4b–c**). These results indicate that centromeric array identity is maintained across haplotypes despite ongoing structural variation.

*Tyba*-array synteny further reveals how centromeric units are redistributed during large-scale genome evolution. In *R. ridleyi*, conserved arrays are repositioned across chromosomes in patterns consistent with rearrangements and fusion events following whole-genome duplication (WGD; **Fig. 3c, Extended Data Fig. 4d**). In *R. pubera*, extensive array synteny is preserved despite two rounds of WGD, with arrays aligning to reconstructed ancestral synteny blocks (**Extended Data Fig. 4e**). Similarly, in the allotetraploid *R. tenerrima*, individual arrays remain traceable across homeologous chromosomes from A and B subgenomes despite extensive post-hybridisation reorganisation (**Extended Data Fig. 4f**). These cases demonstrate that centromeric units can persist through genome duplication and restructuring.

An extreme example of centromeric turnover is provided by *R. filiformis*, which retains near-perfectly fine-scale synteny with *R. tenerrima* yet has lost all but few reduced *Tyba* repeats while preserving the surrounding genomic context (**Fig. 3b, Extended Data Fig. 5**). In several syntenic regions of *R. filiformis*, full *Tyba* arrays present in *R. tenerrima* are reduced to single residual monomers, indicating recent and large-scale erosion of centromeric *Tyba* repeats in *R. filiformis* (**Extended Data Fig. 5c**). This highlights the capacity of holocentromeres to undergo rapid compositional change without compromising chromosome integrity. Quantitatively, arrays unique to individual lineages greatly outnumber those shared between species, even though overall synteny remains high (**Fig. 3d, Supplementary Fig. 6; Supplementary Dataset 3**). In contrast, haplotype-level comparisons show high positional conservation, with variation arising primarily through array copy-number gain or loss (**Fig. 3e, Supplementary Fig. 7; Supplementary Dataset 4**). Together, these patterns indicate that *Tyba* arrays evolve through modular gain and loss events that track with species divergence, whereas intra-species diversity is shaped mainly by fine-scale expansion, contraction, and spacing shifts. Analysis of the genomic context surrounding conserved arrays revealed features associated with long-term array persistence. Between species, but not within species, regions flanking retained array pairs (**Type1/1R**) show a pronounced enrichment of genes within 2–10 kbp of array boundaries (47.50% versus 37.07%, Mann–Whitney U test p = 0.018), a difference that was not detected further from the arrays (10–20 kbp: 38.44% versus 36.97%, p = 0.54) (**Extended Data Fig. 6a–b**). Notably, this pattern was absent at loci where arrays have been lost (**Type2/2R**; 30.42% versus 32.17%, p = 0.689 for 2–10 kbp; 38.44% versus 36.97%, p = 0.644 for 10–20 kbp) (**Extended Data Fig. 6a–d**). This suggests that proximity to transcriptionally active chromatin may contribute to array stability over evolutionary time. In contrast, array boundaries are consistently enriched for short, fragmented TEs, particularly within the first 2 kbp, likely reflecting repeated cycles of array expansion and contraction that disrupt intact elements (**Extended Data Fig. 6a–d**). Consistent with ongoing turnover, sequence identity within *Tyba* arrays is lower than in surrounding regions (81.04% within arrays versus 85.66% in the 15 kbp flanking Type 1/1R regions, Wilcoxon signed-rank test p = 0.002) (**Extended Data Fig. 6a–b**), indicating that even conserved arrays remain highly dynamic at the sequence level.

Together, these analyses show that *Tyba* arrays constitute stable, heritable centromeric units that define a conserved centromeric landscape across haplotypes, accessions, and duplicated genomes. Despite the highly dynamic chromosome architecture of *Rhynchospora*, orthologous *Tyba* arrays persist and maintain centromere identity while undergoing rapid, lineage-specific turnover. This time-dependent balance between stability and diversification underpins the modular and highly plastic nature of repeat-based holocentromeres.

### The TE landscape of repeat-based holocentromeres

We first characterised the genome-wide TE composition across the *Rhynchospora* pangenome. In general, TE abundance was positively correlated with genome size (**Supplementary Fig. 8**). Total TE content contributes between 20% (in *R. ridleyi* and *R. rugosa*) and more than 50% (in *R. pubera* and *R. barbata*) of the genome (**Fig. 2c**). Remarkably, Class I (retrotransposons) and Class II (DNA transposons) share similar abundances in most species (**Supplementary Fig. 9a, Supplementary Dataset 5**). *Ty1/Copia* and *Ty3/Gypsy* LTR retrotransposons are generally found in equal proportions in most species (∼5.7% for both superfamilies for all genomes combined, **Supplementary Dataset 5**). There are four LTR retrotransposon lineages - the *Ty1/Copia Angela* and *Ale* and the *Ty3/Gypsy Athila* and *Ogre* - that are abundant across the *Rhynchospora* genus and collectively occupy 65% of the total lineage fraction of these genomes, with other lineages such as *Ikeros*, *Tekay*, *Retand* and *SIRE* showing moderate and widespread representation, although their relative abundance can vary among species (**Supplementary Fig. 9b, Supplementary Dataset 6**). Class II elements are more homogeneous with *Helitrons* representing the dominant DNA transposon superfamily, followed by *Mutator, hAT*, and *CACTA* with broadly similar proportions across species (**Supplementary Fig. 9c, Supplementary Dataset 5**). Overall, the TE landscape reveals that a shared set of Class I and Class II families is maintained across the genus, despite large differences in total TE content. This broad conservation suggests that a mosaic of transposon types have been retained, reactivated or recurrently amplified throughout *Rhynchospora* evolution.

Next, we characterised the TE content and characteristics within *Tyba*-defined centromeric space (**Fig. 4a–e; Extended Data Fig. 7**). TEs occupied a substantial fraction of these regions with an average of 14.5% across the genus, although their abundance varied considerably among species, for example from 6.1% in *R. holoschoenoides* to 27.9% in *R. barbata* (**Fig. 4a, Extended Data Fig. 7, Supplementary Dataset 5**).

**Fig. 4:**
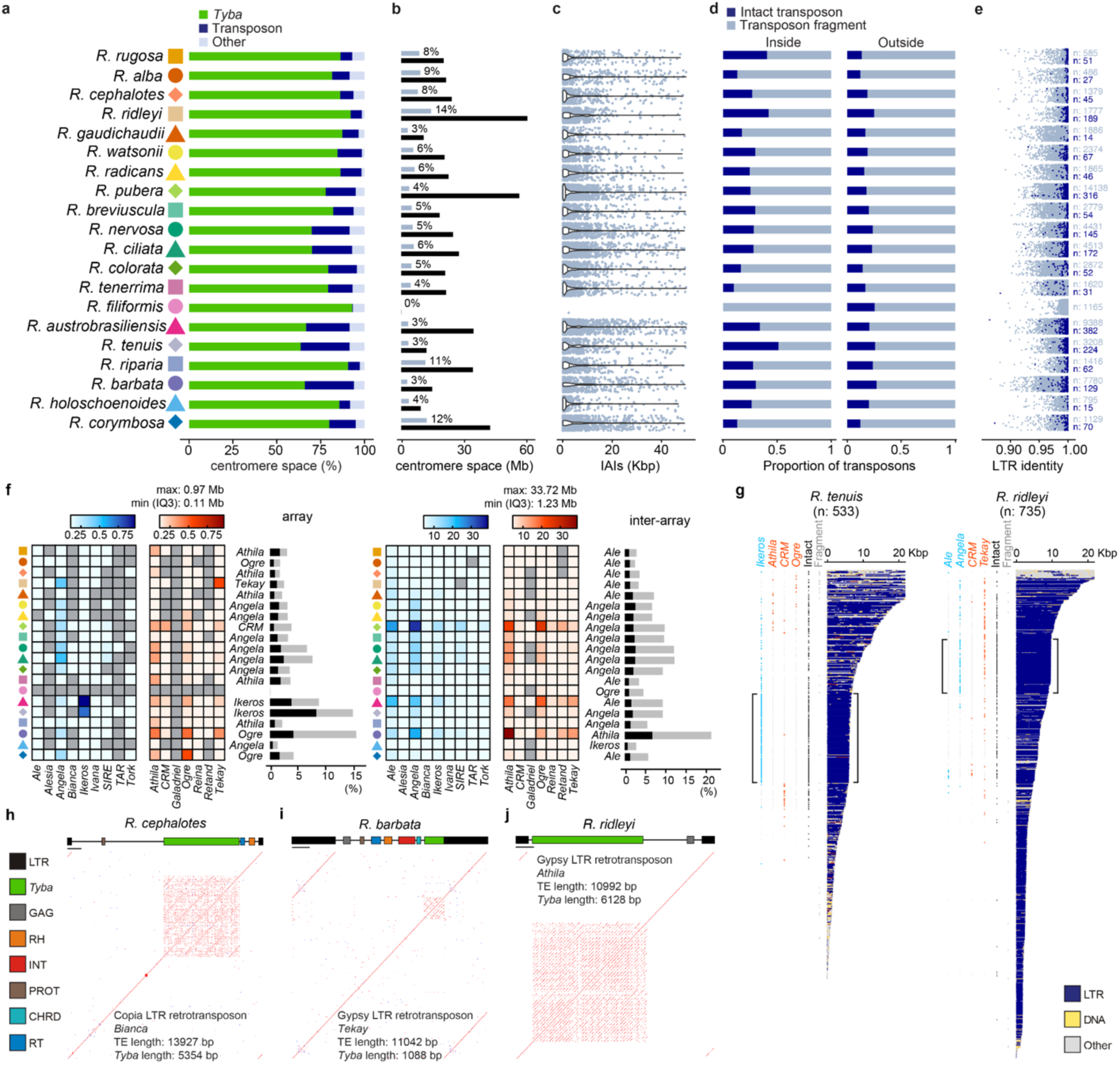
The TE landscape of *Tyba*-based holocentromeres in *Rhynchospora*. (**a–e**) Quantitative characterisation of the centromeric space occupied by TEs across 20 *Rhynchospora* species. (**a**) Bar plots showing the proportion of the centromeric arrays that are composed of *Tyba* repeats (green), transposons (dark blue), or other sequences (light blue). (**b**) Total size of the holocentromere satellite regions in megabases (black bars, in Mbp) and their proportion of the genome (light blue). (**c**) Size distribution of intra-array interruptions (IAIs). The mean size of IAIs is consistent across species, ranging between 2.3 and 8.0 kbp. (**d**) Bar plots showing the proportion of intact (dark blue) versus fragmented (light blue) transposon annotation, inside the centromeric *Tyba* arrays versus outside. (**e**) LTR identity values for intact LTR retrotransposons inside (dark blue), or outside (light blue), the centromeric *Tyba* arrays. (**f**) Patterns of centrophilic LTR retrotransposon lineages. Each plot shows the relative abundance of LTR lineages inserted within *Tyba* arrays (left) compared with those inserted in the rest of the genome (right). Relative abundance was calculated as the proportion of DNA space of every lineage divided by the total DNA space of all LTR lineages combined within (left) and outside (right) the *Tyba* arrays. Copia lineages are shown with blue shading, Gypsy lineages with red shading. Grey cells indicate the complete absence of a lineage. The bar plots show the proportion that all LTR lineages combined occupy (grey bar) within (left) and outside (right) of the *Tyba* arrays, with the black bar showing the contribution of the most abundant lineage in each occasion. Centromeric regions are characterised by a distinct, often enriched subset of LTR lineages relative to the genomic background. (**g**) *Tyba* intra-array interruptions (IAIs) of indicated species are shown, ordered according to their physical length in kilobase pairs (kbp), and coloured according to their transposon content along their sequence. IAI lengths are capped at 20 kbp for visual clarity. The presence of different LTR lineages (Ty1/Copia: blue, Ty3/Gypsy: red) is included on the left of every IAI. Two additional columns indicate intact elements (black) and their fragments (dark grey) to emphasize the large number of IAIs that are composed entirely by a single intact element (indicated by brackets). Data for all species are shown in **Extended Data** Fig. 9. Source data available in **Supplementary Datasets 5 and 6**. (**h–j**) Representative sequence identity dot plots showing examples of *Tyba* insertions into intact LTR retrotransposons with variable degrees of structural preservation and integration. Each schematic illustrates *Tyba* units/sequences (green) embedded within canonical LTR element structures such as Ty1/Copia Bianca in *R. cephalotes* (**h**), Ty3/Gypsy Tekay in *R. barbata* (**i**) and Ty3/Gypsy Athila in *R. ridleyi*. Horizontal bars in **h–j** correspond to 1 kbp. More examples of both intact and fragmented TEs with signs of *Tyba* insertion are shown in **Supplementary** Fig. 11.

LTR retrotransposons accounted for most of the centromere space (11.7% on average), with DNA transposons and unclassified repeats comprising the remainder (**Extended Data Fig. 7**). Intact elements, rather than TE fragments, are more frequent within *Tyba* arrays than in the rest of genomic space (*p* = 0.006292; **Fig. 4d**), while they also show higher LTR identity compared with elements outside the centromeres (**Fig. 4e**). These patterns indicate that centromeric TEs are evolutionarily younger, more recently active, and continuously recycled within the *Tyba* arrays. They also suggest that holocentromeres, despite their diffuse organisation, remain permissive to TE invasion and renewal. A similar enrichment of young retrotransposons has been observed in monocentric centromeres^14,33^, pointing to a shared evolutionary mechanism in which active TEs recurrently reshape satellite-rich DNA and contribute to centromere evolution.

To examine the composition of TEs associated with *Tyba*-based centromeric units, we classified all LTR retrotransposon insertions found within intra-array interruptions (IAIs) of *Tyba* centromeres across 20 *Rhynchospora* species using lineage-resolved annotation (**Fig. 4f, Extended Data Fig. 8, Supplementary Dataset 6**). The analysis revealed a remarkable diversity of “centrophilic” LTR lineages, i.e. elements that are localised within centromeric space. Some lineages such as Ogre in *R. corymbosa* and *R. barbata*, and Tekay in *R. ridleyi* showed species-biased enrichment. Similarly, Ikeros has specifically invaded the centromeres of *R. tenuis* and *R. austrobrasiliensis* that are closely related on the phylogenetic tree. These two species have a large number of IAIs that are made up in their entirety by intact Ikeros elements (73 and 92 respectively), also *R. ridleyi* with intact Tekay and Angela elements (49 and 47 respectively), consistent with very recent invasion of their centromere space (**Fig. 4g, Extended Data Fig. 9, Supplementary Fig. 10**). In contrast, the two most abundant lineages within the centromeres, Angela and Athila that occupy 22.2% and 19.6% respectively of the total fraction of all LTR lineages within the centromeres, were also the most abundant outside the centromeres (19.9% and 17.6%) (**Fig. 4f, Extended Data Fig. 8, Supplementary Dataset 6**). This mirroring of genome-wide insertion dynamics suggests that the scattering of numerous *Tyba* arrays along the chromosomes can create ample opportunities for centromere invasion by multiple TE types simultaneously. Notably, the third most abundant LTR lineage across all *Rhynchospora* genomes, Ale, was largely absent within *Tyba* arrays (2.7% of the total fraction of all LTR lineages compared to 16.1% outside the centromeres), indicating that a higher density of centromeres does not necessarily lead to increased invasion levels (**Fig. 4f, Extended Data Fig. 8, Supplementary Dataset 6**). Taken together, the coexistence of multiple TE lineages within *Tyba*-rich regions suggests that holocentric architecture in *Rhynchospora* provides a broader genomic context for TE colonisation, resulting in a mosaic centromeric landscape shaped by species-specific, TE-specific and stochastic insertion events, contradicting satellite-based monocentric genomes where, typically, a very small number of TEs are centrophilic in a single genome^14,34,35^.

Because we have previously found the occurrence of the Helitron TCR1 element carrying and dispersing *Tyba* arrays in the *R. pubera* genome^9^, we sought to explore the structural interplay between *Tyba* and TEs at a higher sampling of species. Thus, we manually identified several examples of intact elements with embedded *Tyba* insertions as long as 41 kbp from diverse transposon superfamilies and LTR lineages. This includes Ty1/Copia (Bianca), Ty3/Gypsy (Athila and Tekay), and CACTA DNA elements, suggesting multiple independent insertion and recombination events (**Fig. 4h–j, Supplementary Fig. 11a–c**). We also catalogued the remarkable diversity of TE fragments from multiple superfamilies that carry *Tyba* sequences, including LTR and non-LTR retrotransposons, DNA transposons and Helitrons (**Supplementary Fig. 11d**). Overall, our data reveal a broad spectrum of dynamic interplay between satellite DNA and transposons within holocentric *Rhynchospora* chromosomes. *Tyba* arrays are not isolated sequence islands but instead form part of a larger repetitive landscape together with transposons, which jointly shapes and contributes to the structural renewal and long-term evolution of repeat-based holocentromeres in *Rhynchospora*.

### A role for the spatial organisation of *Tyba* arrays in controlling chromatid architecture

To investigate how the spatial organisation of *Tyba* arrays contributes to chromosome architecture, we quantified *Tyba* abundance, array size and inter-array spacing across 20 *Rhynchospora* species. Although gene and repeat diversity follow phylogenetic structure, neither array length nor inter-array spacing shows evidence of phylogenetic constraint, indicating that centromeric organisation evolves rapidly and independently of species relationships (**Fig. 5a–b**).

**Fig. 5:**
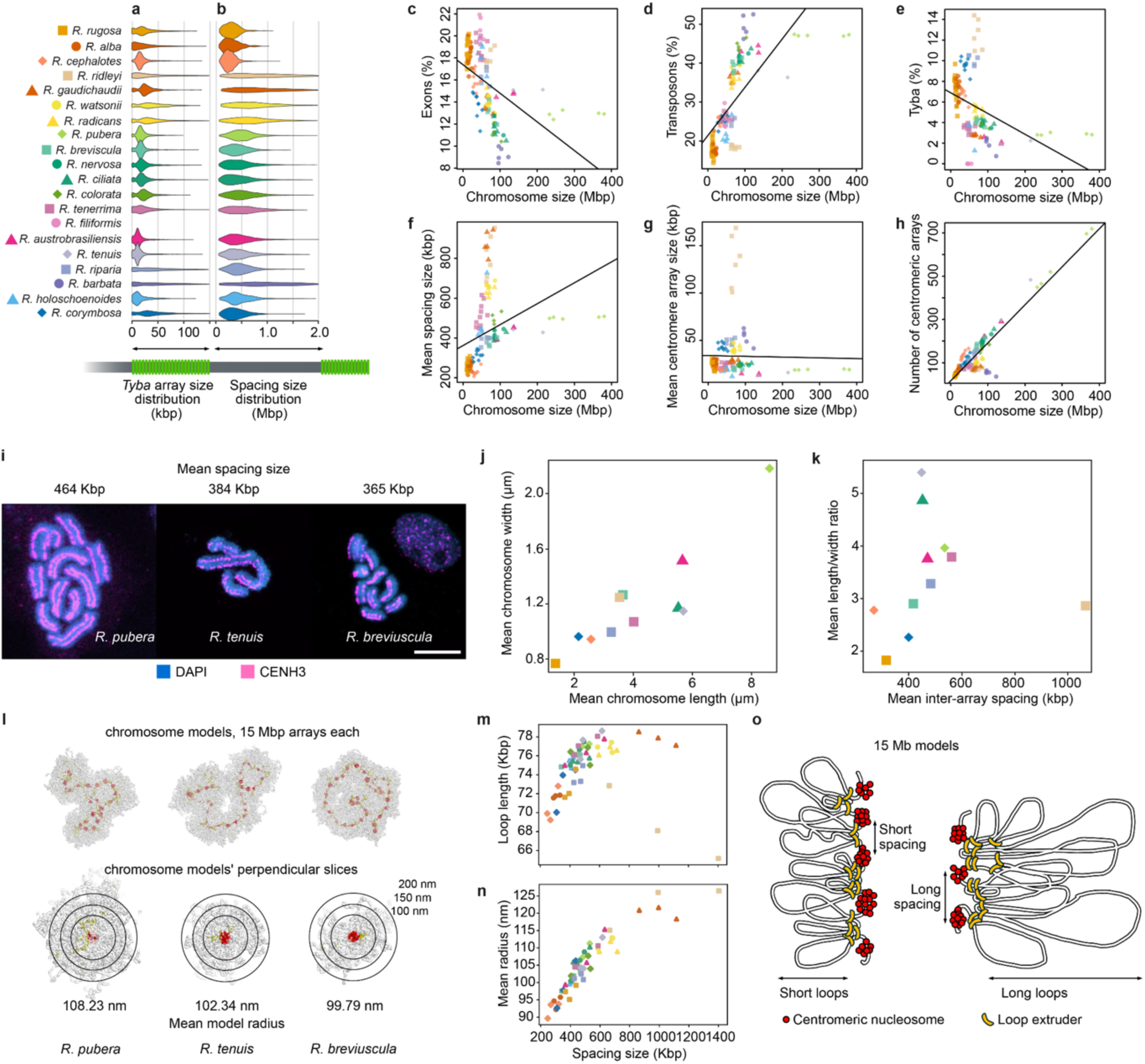
Scaling relationships between *Tyba* arrays and chromosome size control chromatid architecture in *Rhynchospora*. (**a**) Distribution of individual *Tyba* array sizes (kbp) showing stable array lengths across species despite large differences in genome and chromosome size. (**b**) Distribution of inter-array spacing distances (Mbp), revealing that spacing, but not array size, scales with chromosome size. (**c–e**) Variation in total gene (**c**), TE (**d**) and *Tyba* repeat (**e**) abundance across 20 *Rhynchospora* species, expressed as a percentage of the chromosome size. (**c**) Negative correlation between gene content and chromosome size. (**d**) Positive correlation between transposable element (TE) content and chromosome size. (**e**) Negative correlation between *Tyba* content and chromosome size, indicating antagonistic dynamics between *Tyba* and TE repeat classes. (**f**) Average inter-array spacing distances (Mbp) and (**g**) array size (kbp), revealing that spacing, but not array size, scales with chromosome size. (**h**) Relationship between chromosome length and *Tyba* array number, showing near-linear scaling. (**i**) Metaphase chromosomes stained with DAPI (blue) and CENH3 (magenta) of three representative *Rhynchospora* species depicting different chromosome morphologies and *Tyba* centromeric inter-array spacings. (**j**) Cytological quantitative relationship between chromosome length and width across eleven *Rhynchospora* species with varying chromosome morphologies. Chromosomes were stained with DAPI to obtain the measurements. (**k**) Cytological quantitative relationship between chromosome width and mean *Tyba* array inter-spacing across ten *Rhynchospora* species. (**l**) Polymer simulations of holocentric chromosomes with three representative inter-array spacing regimes. Closely spaced *Tyba* arrays produce compact, thin chromatids, whereas wider spacing generates extended chromatin loops and thicker chromatids, perpendicular slices of simulated chromosomes visualising internal chromatin organisation. (**m**) Positive correlation between simulated loop length and inter-array spacing across 18 *Rhynchospora* species. (**n**) Positive correlation between simulated chromatid mean radius length and inter-array spacing across 18 *Rhynchospora* species. (**o**) Model summarising how holocentric chromosome width variation arises through combined changes in array number and inter-array spacing, maintaining chromosome-wide centromere coverage.

Given earlier evidence for antagonistic interactions between *Tyba* and TE content^22^, we compared gene (exons), transposon and *Tyba* abundances as a function of chromosome size (**Fig. 5c–e**). Chromosome lengths vary by an order of magnitude (from 9 Mbp in *R. rugosa* to 375 Mbp in *R. pubera*), yet gene content decreases with size, showing that genome expansion is not driven by gene duplication (Wald t-test, *p* < 0.00005, **Fig. 5c**). By contrast, TE abundance increases markedly with chromosome size (Wald t-test, *p* < 0.00005, **Fig. 5d**), while *Tyba* repeats are more abundant on shorter chromosomes (Wald t-test, *p* < 0.00005, **Fig. 5e**). These opposing trends point to an evolutionary trade-off between *Tyba* accumulation and TE expansion.

A key finding is that inter-array spacing increases with chromosome size (Wald t-test, *p* = 0.00012, **Fig. 5f**), whereas individual array length remains relatively constant (Wald t-test, *p* = 0.67447, **Fig. 5g**). At the same time, the number of arrays scales nearly linearly with chromosome length (Wald t-test, *p* < 0.00005, **Fig. 5h**). Together, these patterns indicate that larger chromosomes accommodate more *Tyba* units and larger intervals between them. This dual mechanism of array addition and spacing expansion is consistent with predictions of holokinetic drive^26^, in which selection during asymmetric meiosis shapes chromosome size through cycles of chromosome fissions with repeat loss or fusions with repeat gain.

Moreover, the variation in inter-array spacing correlates strongly with differences in chromosome morphology. Species with widely spaced arrays, such as *R. pubera*, possess visibly thicker and longer metaphase chromatids, whereas those with narrow spacing, such as *R. breviuscula*, have slender chromosomes (**Fig. 5i and Extended Data Fig. 10a**). Quantitative cytogenetic measurements confirmed links between chromatid width, chromosome length and mean inter-array spacing across the genus (**Fig. 5j–k**).

To determine whether *Tyba* spatial arrangement can mechanistically explain these morphological differences, we carried out enhanced polymer simulations^36^ using empirically measured array and spacing values for 18 species (**Fig. 5l–n**). Chromosome-scale models were built as stretches of 15 Mb of alternating centromeric units (array) and inter-array spacing (**Fig. 5l and Supplementary Dataset 7**). Molecular dynamics simulations incorporated loop extrusion as a compaction mechanism for inter-array spacing regions, which brings centromeric arrays to a line (**Supplementary Movies 1 and 2**). The compaction of three different stretches of each of 18 *Rhynchospora* species illustrates a progressive thickening of chromatids as spacing increases, a trend recapitulated in perpendicular cross-sections of polymer models (**Fig. 5l, Extended Data Fig. 10b**). Consistent with an anchoring role for centromeric units in loop extrusion, wider spacing reduced the frequency of anchoring events, permitting larger loops and yielding thicker chromatids. Accordingly, loop-length distributions shifted toward larger values as spacing increased, and the mean chromatid radius rose from 89.7 nm at 247 kbp spacing of *R. cephalotes* to 126.5 nm at 1.4 Mbp of *R. ridleyi* (**Fig. 5m–n and Supplementary Fig. 12**).

These combined cytogenetic and modelling results show that the organisation of *Tyba* arrays governs higher-order chromatin folding in holocentric chromosomes. Inter-array spacing potentially acts as a tunable parameter controlling loop size and, consequently, chromatid thickness. This spacing-dependent biophysical scaling provides a mechanistic basis for the striking morphological diversity observed across *Rhynchospora* and offers a means by which dispersed centromeric units maintain balanced kinetochore engagement during holocentric chromosome segregation. Our findings further support the earlier proposal that loop extrusion mechanisms are modulated by centromeric units^36^.

## Discussion

Our pan-genome analysis reveals a unifying and scalable logic for holocentromere organisation in *Rhynchospora*, centred on the deeply conserved satellite repeat *Tyba*. Despite more than 40 million years of divergence, *Tyba* remains the single centromeric repeat family across the genus, demonstrating an exceptional degree of evolutionary stability relative to the rapid satellite replacement common among monocentric taxa^13,14,37^. Within this conserved molecular framework, *Tyba* arrays behave as modular units that are dynamically gained, lost, split and merged across species and haplotypes, yet collectively maintain chromosome-wide centromere coverage. This duality between deep sequence conservation and intense local structural turnover underpins the distinct evolutionary trajectory of repeat-based holocentromeres.

A key discovery is that holocentromeres scale not by enlarging individual arrays but by modulating the number and spacing of *Tyba* units as chromosomes expand or contract. Array size remains remarkably constant across the genus, whereas both array count and inter-array spacing increase with chromosome length. This architectural scaling aligns with predictions of holokinetic drive^26^, whereby chromosome size biases kinetochore activity during asymmetric meiosis, favouring cycles of fission or fusion that respectively reduce or increase repeat content. Our synteny-guided tracking shows that these cycles are executed through the modular reorganisation of centromeric units, providing a mechanistic path by which chromosome number and size evolve without compromising centromere function.

Transposable elements further contribute to this dynamic, with centromeric space enriched for specific LTR retrotransposon lineages that preferentially or stochastically by proximity insert into *Tyba* arrays. These TE-satellite interactions create opportunities for sequence renewal while preserving the overarching identity of the centromeric repeat family. Together, the stable repeat identity and modular organisation of arrays explain how holocentromeres withstand extensive karyotype changes while maintaining stable chromosome segregation. Consistent with this view, comparisons of syntenic array pairs between species, representing deeper evolutionary divergence than haplotype comparisons, reveal a marked enrichment of gene density in regions flanking retained arrays, a feature absent from loci where arrays have been lost, suggesting that transcriptionally active chromatin may contribute to the long-term persistence of centromeric units.

Our work also uncovers a direct mechanical role for holocentromere architecture in shaping chromosome organisation. Polymer simulations and cytogenetic analyses show that the spacing between *Tyba* units governs chromatin loop length and chromatid thickness, with wider spacing producing longer loops and thicker chromatids. This places *Tyba* inter-array spacing within the broader framework of chromosome biology, where chromosome size determines loop size and axis compaction. Previous work has shown that smaller chromosomes form shorter loops and longer axes per megabase, promoting higher crossover density, whereas larger chromosomes with longer loops and compressed axes experience lower recombination^38–40^. Our findings provide the mechanistic bridge between these principles: by modulating loop length, *Tyba* spacing translates chromosome size into loop geometry. This interpretation is reinforced by our Hi-C–based analyses and our *Rhynchospora* recombination study, which show that chromosome size predicts recombination rate through chromatin and axis geometry^6^. Together, these results position *Tyba* repeats as a key architectural parameter linking holocentromere organisation, chromosome size and the quantitative reshaping of recombination landscapes in holocentric *Rhynchospora*.

Finally, the profound conservation of *Tyba* in *Rhynchospora* parallels the persistence of holocentromeric satellites in *Luzula*^20^, *Carex*, *Schoenoplectus* and other holocentric plant lineages^33^, suggesting that slow turnover of the core centromeric repeat may be a general feature of repeat-based holocentromere evolution. In contrast with monocentric species such as *Arabidopsis*, where centromeric satellites are frequently replaced over short evolutionary timescales^14^, *Rhynchospora* appear to evolve primarily through structural reorganisation of an otherwise stable *Tyba* repeat. This emerging paradigm reframes how we understand centromere identity as holocentric chromosomes evolve less by sequence invention and more by scalable reconfiguration of conserved modules.

Together, our findings establish a mechanistic framework linking centromere architecture, chromosome mechanics and genome evolution. They highlight how repeat-based holocentromeres can combine deep molecular continuity with remarkable structural flexibility, offering insights into the evolution of genome organisation across eukaryotes and providing a foundation for future studies of centromere dynamics, chromatin physics and meiotic regulation in holocentric systems.

## Methods

### Genome assemblies

All genome assemblies used in the present study were obtained from Zhang et al.^6^ and Zhang et al.^7^.

### Tandem repeat, transposable element and gene annotation

We annotated centromeric satellite repeats and TEs across 56 chromosome-level haplotype assemblies from 20 *Rhynchospora* species. Tandem repeat arrays were identified using TRASH2^41^, which detects and quantifies satellite DNA families based on periodicity and sequence similarity. TEs at the superfamily level were annotated with EDTA^42,43^ (--anno 1, --force 1). We filtered the initial annotation by intersecting TE coordinates with satellite repeat coordinates from TRASH2. TEs that overlapped by >80% of their length with satellites were removed. LTR retrotransposons were classified into lineages using TEsorter^44^ (-db rexdb-plant - nolib). Elements showing inconsistent superfamily classification between EDTA and TEsorter were removed, as were those with no functional domains or unknown Clade/Lineage classification. Gene models were predicted de novo using Helixer^45^ with default parameters (and --lineage land_plant), a deep learning-based tool for plant genome annotation.

### CENH3 ChIP-seq analysis

ChIP was performed following Reimer and Turck^46^ with minor modifications. Young leaves and flowers were collected from the greenhouse grown plants (*R. rugosa*, *R. alba*, *R. ridleyi*, *R. cephalotes*, *R. holoschoenoides, R. filiformis*) and snap frozen. Plant material was fixed by vacuum infiltration in 4% formaldehyde on ice for 1 hour. Nuclei were extracted using NIB buffer (50mM HEPES pH7.4, 5mM MgCl2, 25mM NaCl, 5% sucrose, 30% glycerol, 0.25% Triton X-100, 0.1 % β-mercaptoethanol and 0.1% protease inhibitor) and resuspended in TE-SDS.

Sonication was performed for 30 s ON, 30 s OFF cycles, low voltage with the number of cycles optimised for each plant species as follows: 25 cycles for *R. pubera*, 25 cycles for *R. breviuscula*, 35 cycles for *R. tenuis*, 30 cycles for *R. rugosa*, 30 cycles for *R. alba*, 30 cycles for *R. ridleyi*, 30 cycles for *R. cephalotes*, 30 cycles for *R. holoschoenoides* and 30 cycles for *R. filiformis*. The sonicated chromatin was immunoprecipitated using anti-CENH3 antibody^8^. Antibody-bound chromatin was captured using rProtein A Sepharose® Fast Flow beads (Sigma, GE17-1279-01). The bound chromatin was eluted, de-crosslinked and sent for sequencing. CENH3-ChIP-seq data from *R. pubera*, *R. breviuscula* and *R. tenuis* were retrieved from previous research^7,9,21^.

Raw reads were aligned to the respective genome assemblies using Bowtie2 (v2.5.1)^47^ with default parameters. Peaks were called using MACS3 (v3.0.0a7)^48^ with --broad and --broad-cutoff 0.1 to reflect the diffuse nature of CENH3 binding in holocentrics. Signal coverage, enrichment heatmaps, and correlation analyses were generated with deepTools2 (v3.5.1)^49^, and visualisation was performed in IGV^50^ and pyGenomeTracks^51^.

### Centromeric array identification and higher-order repeat (HOR) scoring

*Tyba* monomers were identified using custom scripts based on repeat periodicity and monomer similarity. Arrays were defined as clusters of ≥5 adjacent *Tyba* monomers with ≤1 kb spacing. Higher-order repeats (HORs) were detected using pairwise alignment of adjacent monomers followed by hierarchical clustering and sliding-window consensus scoring. HOR accumulation was quantified across each array and normalised by array length.

### Synteny analysis of centromeric Tyba arrays

The algorithm performs pairwise comparisons between all identified arrays, considering similarity between upstream and downstream regions, in all strand configurations. Synteny tracking based on repeats alone was tested with multiple methods (sequence alignments, Levenshtein distance, k-mer space comparisons), but proved unreliable due to the rapid turnover of repeat sequence. Instead, 5 kbp long sequences starting at a 2.5 kbp distance upstream and downstream of each array were extracted and queried against whole chromosomal sequences with minimap2 (-x asm20 - k15 -w5 --max-chain-skip 80 --min-occ-floor 20). The 2.5 kbp distance from the array boundary was used to avoid an identified abundance of short TE fragments in that region which initially caused multiple false-positive results. A potential match was made if the distance between hits coming from upstream and downstream sequence mapping was smaller than 500 kbp and both hits were in the same orientation. Since multiple alignment hits were possible, the potential matches were filtered to mirror the original query array as closely as possible. If a match was made to a sequence with a *Tyba* array, it was denoted as Type 1, Type 2 in absence of *Tyba* repeats. Downstream analysis of novel transposon insertions/deletions, repeat expansions/contractions, or array splitting or removal can inform on the evolution of centromeric repeats at a scale not previously reported. For example, two arrays from one of the compared chromosomes can both match with only one array of the other chromosome, and their upstream and downstream similarity levels can indicate that either they used to be together and separated on the first chromosome, or they used to be separate and merged on the other chromosome. This analysis is closely connected to the chromosome size dynamics, as it has been reported that chromosomal fusions and fissions often occur in the centromere-proximal regions in *Rhynchospora*^6,9^.

### Characterisation of TEs associated with Tyba arrays

We analysed in detail the degree and characteristics of TE invasion in the *Tyba*-defined centromeric space. First, within the boundaries of every *Tyba* array, we identified and retrieved all non-*Tyba* sequence shorter than 50 kb, termed Intra Array Interruptions (IAIs). We then quantified and mapped the TE content within IAIs by intersecting the EDTA/TEsorter annotation, allowing us to visualize and analyse the TE profile of IAIs across species (e.g. **Fig. 4g**). Intact and fragmented TEs were profiled separately to explore dynamics of TE insertion and deletion inside versus outside of centromeres, and also to identify IAIs that were generated by recent integration events (i.e. when the length of IAI matches the length of an intact TE). LTR retrotransposons made up the majority of IAIs, however it is expected that some annotations may be missed or misclassified as it is typical with any TE annotation pipeline. Therefore, to increase the resolution and identification at the LTR lineage level, we ran TEsorter over the IAIs sequences themselves and added the new LTR lineage annotations in our analyses. Finally, to identify high-quality cases of *Tyba* repeats within intact TEs, we computationally removed all *Tyba* annotation from the genome assemblies, patched the cutting points, and ran EDTA as in the original genomes. Newly-identified intact elements (that could not be identified previously as intact due to, for example, their much longer length) were annotated for their TE gene content with TEsorter and for the presence and structure of the *Tyba* monomers using re-DOT-able (https://www.bioinformatics.babraham.ac.uk/projects/redotable/).

Alternatively, for the characterisation of *Tyba* fragments found in non-intact TEs (classes I and II) in **Supplementary Fig. 10d**, we applied the Domain-based Annotation of Transposable Elements (DANTE)^52^ pipeline (https://github.com/kavonrtep/dante) and the REXdb database (Viridiplantae_version_3.0). The RT-LTR gene domains identified by DANTE were then used to characterise the LTRs, PPTs, TSDs and PBSs in DANTE_LTR^52^ (https://github.com/kavonrtep/dante_ltr) with the default parameters applied. Following identification, the elements underwent manual curation and annotation to verify the internal structure of the LTRs, based on CDD/NCBI. For the analysis of restricted sections of the pseudochromosomes, the positions of each element were entered into the SRplot tool^53^ and sequences containing both TEs and *Tyba* sequences were extracted using EMBOSS scripts. The sequences of each TE containing *Tyba* insertions were annotated graphically with the IBS-Illustrator tool^54^, using the coordinates of each target sequence.

### Phylogenetic analyses

*Tyba* sequence evolution was assessed using multiple approaches. Principal component analysis (PCA) of representative *Tyba* monomers was performed using the R *prcomp* function on the variances of positional base frequencies in a mafft alignment (--retree 2) . A Levenshtein distance matrix was used to reconstruct *Tyba*-based phylogenetic relationships. Neighbour-joining trees were constructed using FastME^55^, and tree topologies were compared to species phylogenies based on single-copy orthologs.

### Synteny and centromere dynamics

Orthologous gene clusters and synteny blocks were inferred using GENESPACE^56^, which enables comparative genome alignment across multiple species. To track *Tyba* array turnover, we developed a custom synteny-aware array matching algorithm that integrates sequence similarity and flanking genomic context to identify orthologous arrays, including gain, loss, and split-merge events.

### Orthogroup identification and kinetochore protein annotation

Orthologous gene groups were identified using OrthoFinder v2.5.4^57^ across 34 *Rhynchospora* proteomes predicted with Helixer. The dataset included both haplotype-resolved assemblies and haploid assemblies, representing 20 *Rhynchospora* species. OrthoFinder was run with 12 threads (-t 12). The analysis was performed exclusively on *Rhynchospora* proteomes to identify within-genus orthogroups.

A curated set of kinetochore proteins from Arabidopsis thaliana and other plant species was compiled from published datasets^33,58,59^ and used as a reference database for functional annotation. All concatenated *Rhynchospora* proteomes (2,072,986 sequences) were queried against this curated database (55,926 proteins of different eukaryotic species representing 113 proteins of interest) using DIAMOND v2.1.11^60^: *diamond blastp --ultra-sensitive --evalue 1e-5 --iterate --threads 8 --max-target-seqs 50*.

DIAMOND hits were filtered to retain only alignments with ≥25% sequence identity and ≥20% query coverage to account for remote homologs and well-known kinetochore proteins with a high proportion intrinsically disordered regions that show little similarity at the seuqnece level (e.g., KNL2 or CENP-C). Each Rhynchospora protein identified by DIAMOND was then mapped to its corresponding OrthoFinder orthogroup, and the functional annotation (kinetochore protein name) was transferred from the query to the orthogroup level. For each reference kinetochore protein, the best-scoring DIAMOND hit (by bitscore) was selected and its orthogroup assignment was retained. This approach integrates sequence similarity-based homology detection (DIAMOND) with phylogenetically informed orthology inference (OrthoFinder), and all downstream analyses were conducted at the orthogroup level. Reference kinetochore proteins that failed to meet the filtering criteria or lacked orthogroup assignments in Rhynchospora were recorded as unmapped.

### Phylogenetic profiling of kinetochore homologs

For visualization in a phylogenetic profiling matrix, only orthogroups with genes present in at least 20% of Rhynchospora species were included to focus on conserved kinetochore components and reduce noise from lineage-specific genes and misannotations. In the heatmap, orthogroups containing at least one protein with a direct DIAMOND match (e-value < 1×10⁻⁵) to a curated Arabidopsis kinetochore protein were marked with a red star to distinguish validated kinetochore components from putative homologs identified solely through orthogroup membership.

To quantify sequence conservation within each orthogroup, Shannon entropy was calculated for protein multiple sequence alignments. Prior to entropy calculation, alignments were trimmed to remove positions with >80% gap characters. For each remaining alignment position, Shannon entropy (H) was calculated based on Merchant et al.^61^ as follows:

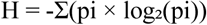

where pi is the frequency of amino acid i at that position (excluding gaps). Entropy values were normalized by dividing by log₂(20) = 4.322, the maximum possible entropy for 20 standard amino acids, yielding normalized entropy values ranging from 0 (complete conservation) to 1 (maximum variability). The mean normalized Shannon entropy was calculated across all positions in each trimmed alignment to provide a single conservation metric per orthogroup.

### Phylogenetic profiling of kinetochore homologs

For CENH3, CENPC, and KNL2 genes, protein sequences from all Rhynchospora species were manually inspected and curated to remove any misannotated homologs. For CENH3, the *Carex depauperata*^33^ ortholog was included as an outgroup. For KNL2, additional sequences were obtained from Zuo et al.^29^ to provide broader phylogenetic context.

Curated sequences were aligned using MAFFT v7.525^62^ with automatic algorithm selection (--auto parameter), and alignments were trimmed using ClipKIT v2.3.0^63^ (--smart-gap). Maximum-likelihood phylogenetic trees were constructed using IQ-TREE v2.3.4^64^. The best-fit amino acid substitution model was selected automatically using ModelFinder^65^ (-m MFP), and branch support was assessed with 1,000 ultrafast bootstrap replicates^66^ (-B 1000). Trees were visualized, annotated, and inspected using iTOL v7^67^ and TreeProfiler^68^ to display gene presence/absence patterns, copy number variation, and other phylogenomic features across the Rhynchospora phylogeny.

### ModDotPlot analysis of concatenated Tyba arrays

Structural analysis of concatenated tandem repeat arrays were performed with ModDotPlot (v.0.9.8)^69^ using the default parameters. ModDotPlot is a dot plot visualization tool designed for analysing the similarity profile of satellite DNA arrays in large sequences and whole genomes. The method outputs an identity heat map by rapidly approximating the average nucleotide identity between pairwise combinations of genomic intervals.

### Slide preparation and CENH3 immunostaining

Slide preparation and immunocytochemistry was performed as described in Castellani et al.^21^. Young flowers of *R. pubera*, *R. tenuis* and *R. breviuscula* were fixated in an ice cold solution of 4% PFA (paraformaldehyde) in PBS (phosphate buffered saline solution, pH 7.5, 1.3 M NaCl, 70 mM Na2HPO4, 30 mM NaH2PO4) with 0.1% Tween-20 for 30min in vacuum. Afterwards, flowers were dissected on a glass slide and only very young anthers were isolated. These anthers are known to contain mitotic stages prior to the initiation of meiosis. Mitotic nuclei were released in a drop of 1XPBS with 0.1% Triton x-100, stirred and squashed with a coverslip, later removed with liquid nitrogen. Samples were mounted with a solution of DAPI 0.2 µg/ml in Vectashield and checked for mitotic stages. Specimens were permeabilised and blocked with a solution of 3% BSA (bovine serum albumin) in PBS with 0.1% Triton x-100.

Antibody from Marques et al.^8^, raised in rabbit against RpCENH3 (peptide ARTKHFSVRSGKKSASRTK) was diluted 1:200 (v:v) in the same blocking buffer and incubated with the samples overnight at 4°C. On the following day slides were washed in PBS+0.1% Triton x-100 3 times for 5min each. Slides were then incubated for 2h at room temperature with an anti-rabbit secondary antibody raised in goat conjugated with STAR ORANGE (Abberior, STORANGE-1007). Washes were repeated as with the primary antibody and slides were mounted again in Vectashield with DAPI 0.2 µg/ml. Images were acquired with a Zeiss Axio Imager Z2 with Apotome system for optical sectioning and deconvolution. Images were processed in ZEN and Adobe Photoshop.

### Chromosome measurements

Mitotic chromosomes were prepared from root tip meristems of nine additional species (*R. austrobrasiliensis, R. ciliata R. corymbosa, R. filiformis, R ridleyi, R. riparia, R. rugosa, R. tenerrima, R. cephalotes*) to enable the measurements of chromosome length and width. Briefly, root tips were fixed in a 3:1 (v/v) ethanol:glacial acetic acid solution for one day at room temperature followed by an additional day at 4°C. The material was then washed in ice-cold water and digested in a solution containing 4% cellulase (Onozuka R10), 2% pectinase in 0.01M citrate buffer (pH 4.5) for 60 - 90 minutes at 37°C. After After digestion, the samples were washed three times in citrate buffer, once in ethanol, and finally suspended in a 1:3 (v/v) ethanol:glacial acetic acid solution. The suspension was dropped onto a slide maintained at 50 °C and high humidity for 2 min to promote the spreading of metaphase chromosomes and finally dried for 1 minute at 50 °C without humidity as described in et al.^70^. Images were acquired as previously described and measurements of length and width were done using ZEN software utilities.

### Polymer simulations

Polymer modelling was used to simulate the mechanical impact of centromeric array organisation on chromosome condensation following the previous method from Câmara et al. ^36^ with modifications (**Supplementary Dataset 7**). Chromosomes were modelled as beads-on-a-string polymers, where each nucleosome is one bead (∼200 bp), using empirical *Tyba* array size and inter-array spacing values. Simulations were performed with OpenMM^71^ and analysed with custom Python scripts (https://github.com/InsilicoGenebankProteomics/Rhynchosp <u>ora_width</u>). Chromatin loop length and chromatid thickness were measured across varying inter-array distances to predict mechanical consequences of centromeric modularity, and validated against cytological data.

## Data availability

ChIP raw sequencing data have been deposited in the European Nucleotide Archive under project PRJEB105456. Custom scripts are available at https://github.com/vlothec/Rhynchospora. Genome assemblies are available from Zhang et al.^6,7^. The annotations presented in this work are made available for download at EDMOND, the Open Research Data Repository of the Max Planck Society. All other data are available from the corresponding author upon request.

## Contributions

A.M. and P.W. conceived the research and supervised the project. P.W. performed tandem repeat analysis, repeat synteny tracking. E.P.R., L.A.R. and A.B. performed transposable elements analysis. A.M. performed ModDotPlot analysis. A.S.C. performed the polymer modelling of loop architecture. G.T. and M.C. performed the immunostaining experiments. L.A.R. performed cytology and chromosome measurements. G.T. and M.Z. performed the ChIP–seq analysis. L.M.P. and A.L.L.V. performed repeat analysis.

B.H. performed ChIP–seq libraries. J.G and I.H. performed kinetochore genes’ analysis. A.M. wrote the first manuscript draft with input from all authors. All authors approved the final version of the manuscript.

## Acknowledgements

This study was funded by the Max Planck Society (core funding to A.M.), the German Research Foundation (MA 9363/2-1 and MA 9363/3-1), and the European Union (European Research Council Starting Grant, HoloRECOMB, grant no. 101114879 to A.M.). The DFG also funded this work under Germany’s Excellence Strategy—EXC 493 2048/1–390686111 (to A.M.). M.Z. and A.S.C are financially supported by the DFG (grant no. MA 9363/2-1 and SO 2132/1-1, respectively). Research was supported by ERC grant 101142254 EvoPanCen to I.H. and a “la Caixa” Fellowship LCF/BQ/EU24/12060051 to J.G.I. This study was supported by Royal Society awards UF160222, RF/ERE/221032, 1004 URF/R/221024, RGF/R1/180006, RGF/EA/201030, and RF/ERE/210069 to A.B. This work was supported by the National Science Center grant 2024/52/C/NZ2/00246 to P.W.

---

**Extended Data Fig. 1.**
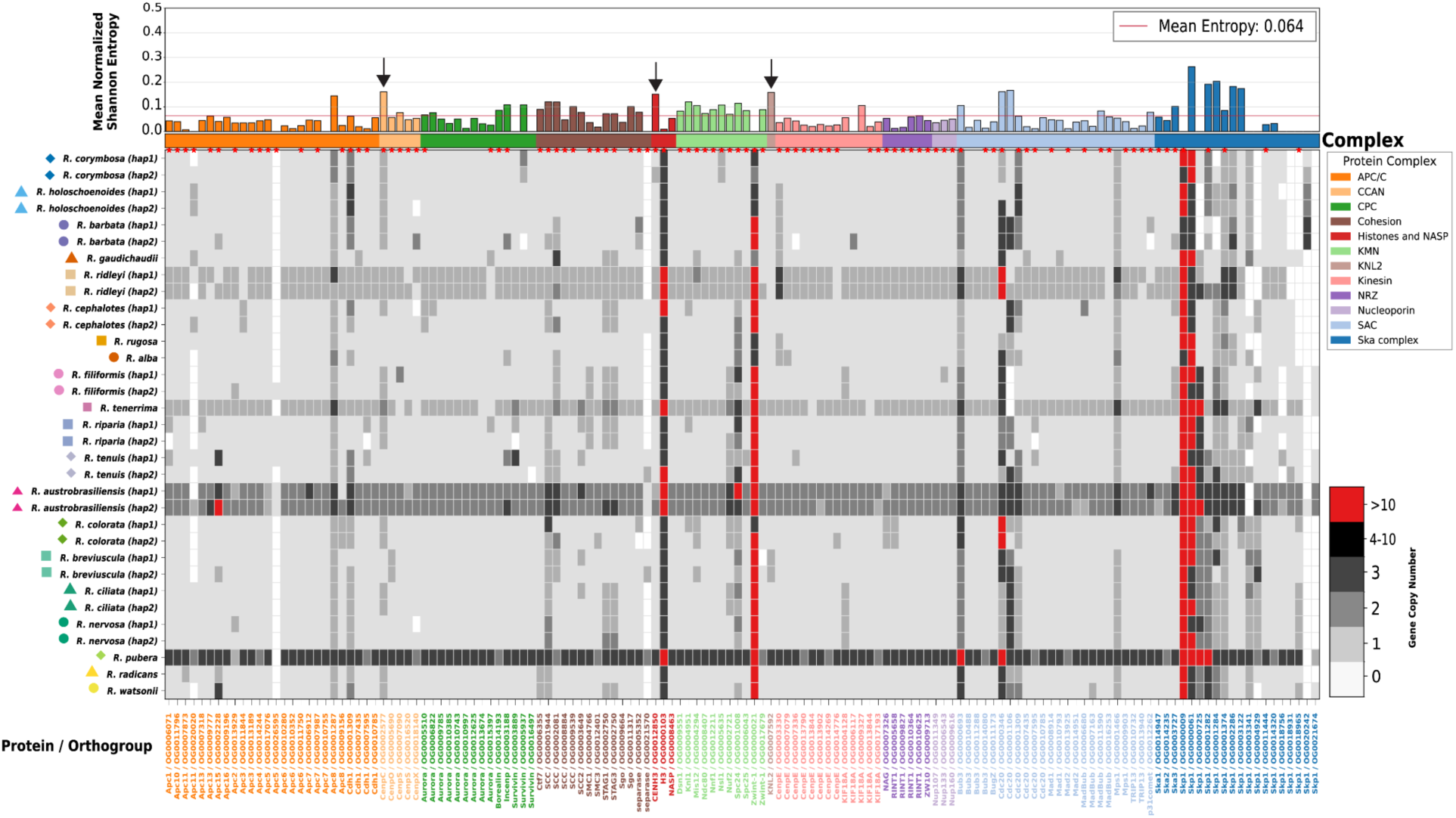
: Landscape of kinetochore proteins in *Rhynchospora*. Phylogenetic profile of kinetochore (and related complexes in cell division) protein orthogroups across *Rhynchospora* species. The entropy plot (top) shows mean normalized Shannon entropy values (0-1) indicating sequence conservation within each orthogroup, with lower values representing higher conservation. Red stars indicate orthogroups with direct matches to *Arabidopsis* kinetochore proteins. Heat intensity reflects gene copy number. Arrows indicate CENP-C, CENH3 and KNL2, showing high entropy rates well above the mean, suggesting rapid evolution.

**Extended data Fig. 2:**
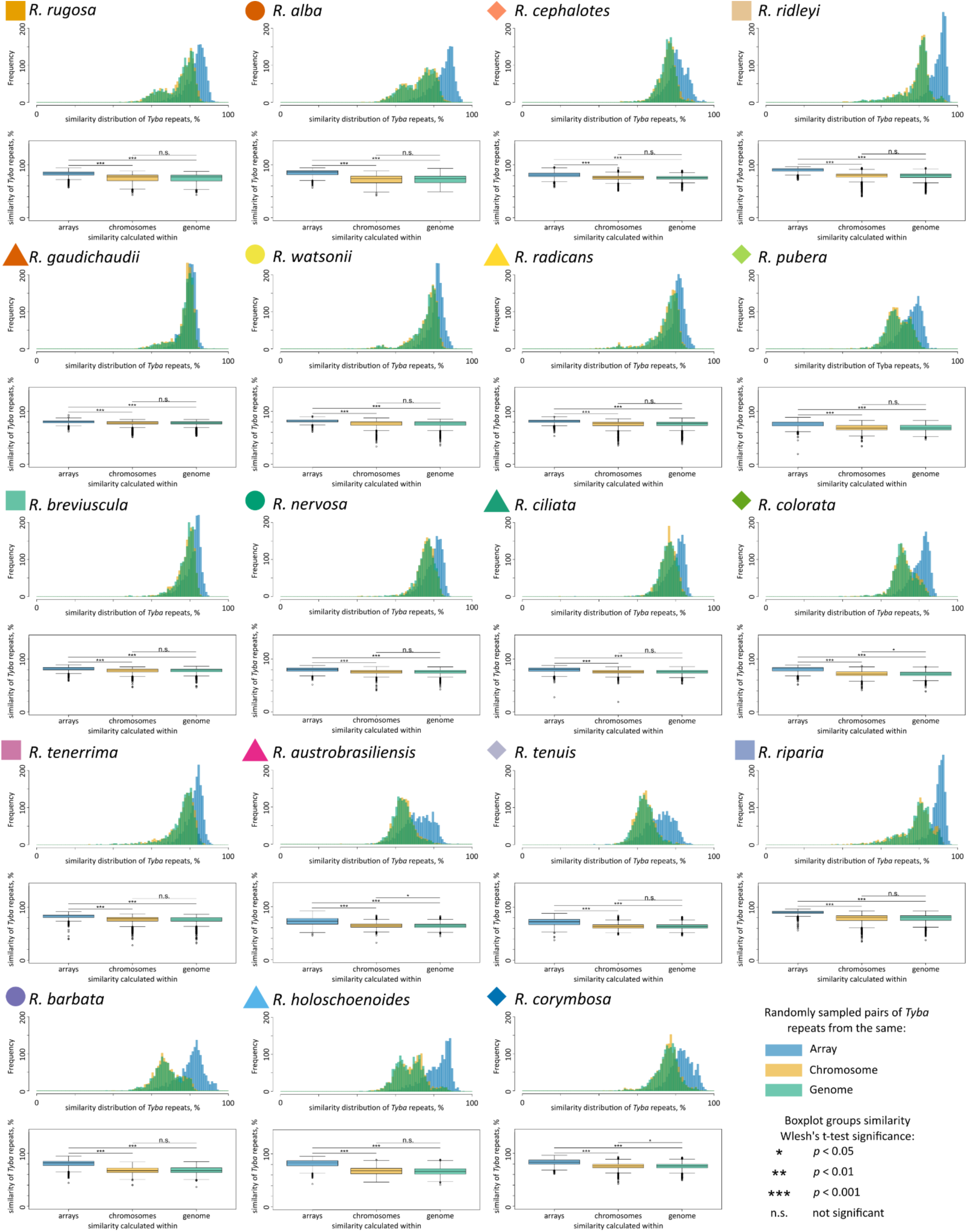
Hierarchical similarity of *Tyba* monomers across arrays, chromosomes and genomes in *Rhynchospora*. Similarity distributions of *Tyba* monomers calculated at three hierarchical levels: individual arrays (blue), chromosome-wide (yellow) and genome-wide (green) for 20 *Rhynchospora* species. For each species, histograms show the distribution of pairwise sequence similarity between randomly sampled *Tyba* monomer pairs drawn from the same array, chromosome or genome, while boxplots summarise the corresponding similarity distributions. Across species, *Tyba* repeats exhibit the highest similarity within individual arrays and similar reduced similarity at the chromosome and the genome-wide levels, consistent with strong local homogenisation and progressive diversification at broader genomic scales. Statistical significance between levels was assessed using Welch’s t-tests (*p* < 0.05, p < 0.01, *p* < 0.001; n.s., not significant).

**Extended Data Fig. 3:**
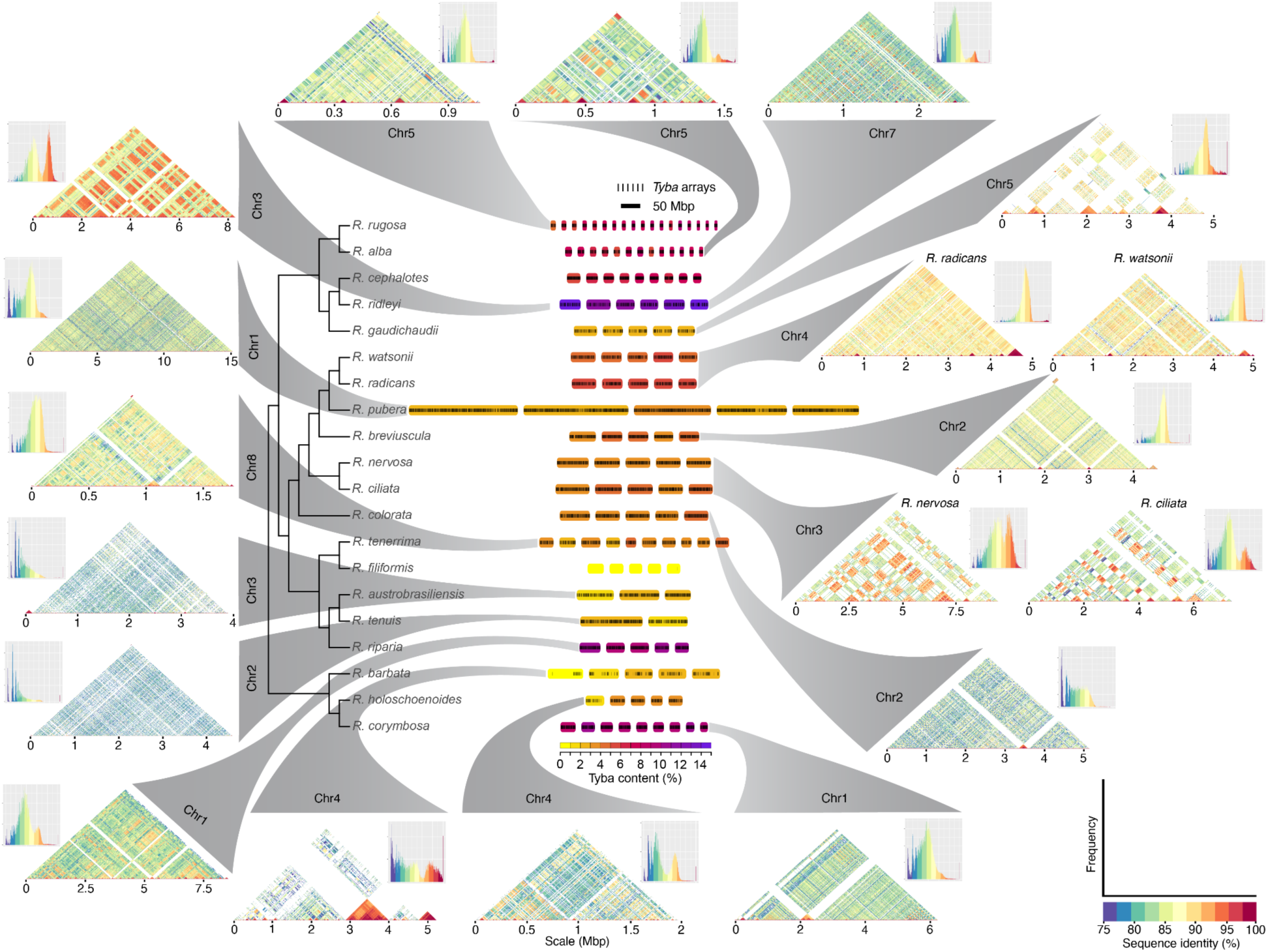
Chromosome-scale organisation, similarity and phylogenetic distribution of *Tyba* holocentromeric arrays across the *Rhynchospora* pangenome. Chromosome-scale maps summarising the distribution, abundance and sequence similarity of *Tyba* arrays across 20 *Rhynchospora* species, organised according to the species phylogeny (center). For each species, coloured blocks indicate the genomic abundances of *Tyba* arrays along individual chromosomes, with colour intensity reflecting relative *Tyba* content. *Tyba* array distribution is represented by black bars. Grey ribbons connect example chromosomes for each species, illustrating the diverse levels of intrachromosomal homogenisation of satellite repeat arrays despite conserved synteny. For representative chromosomes, triangular heatmaps show pairwise sequence similarity among *Tyba* repeats along the chromosome, with accompanying histograms indicating similarity distributions. Together, the figure highlights the conserved yet flexible organisation of *Tyba* holocentromeres, revealing stable chromosome-wide coverage alongside pronounced interspecific variation in array number, spacing and internal similarity.

**Extended Data Fig. 4.**
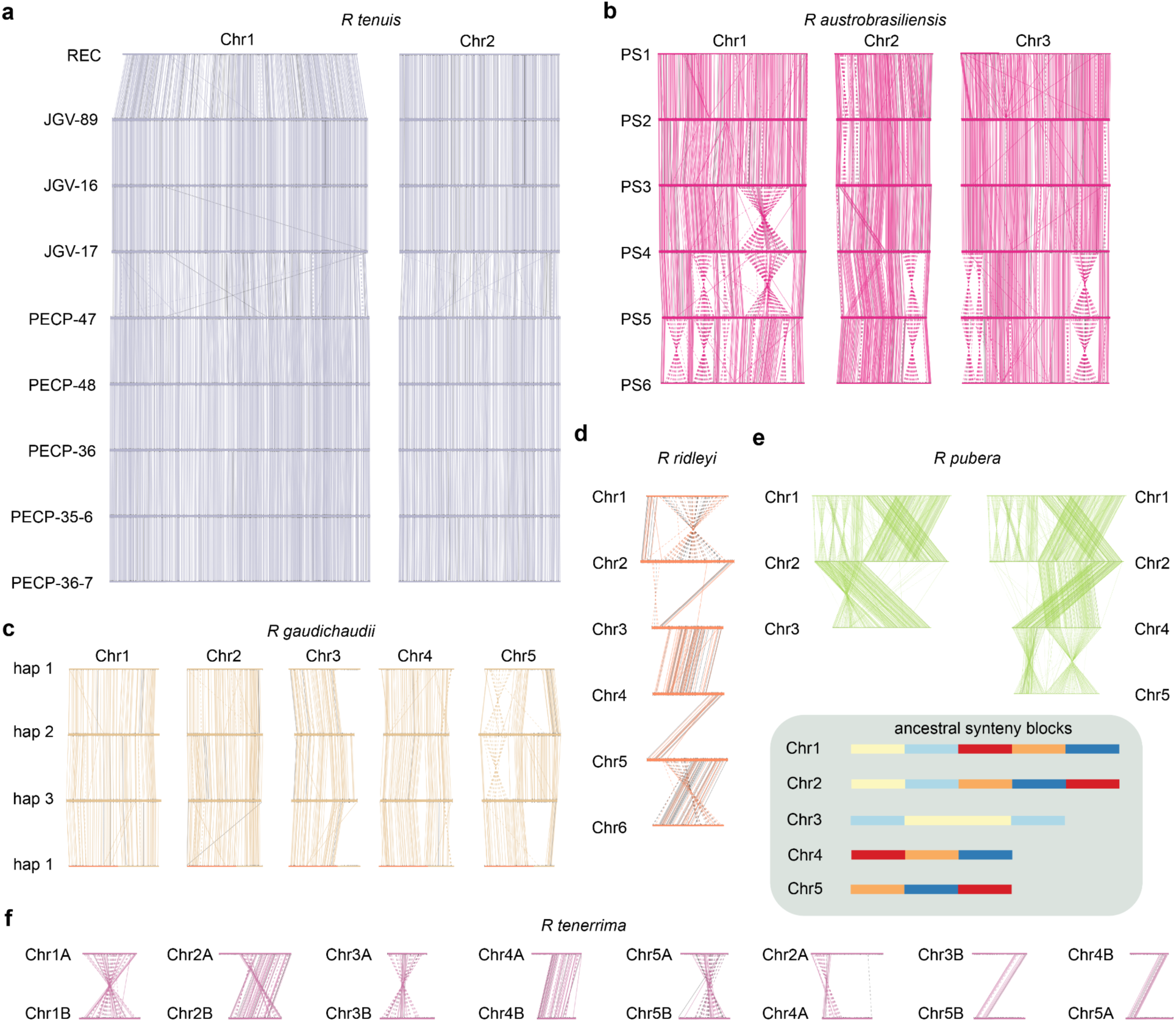
: *Tyba*-array synteny across haplotypes and duplicated genomes reveals conserved centromeric landscape despite karyotype restructuring. *Tyba*-array synteny: (a) for chromosomes 1 and 2 from haplotype 1 across of nine *R. tenuis* accessions; (b) among phased pseudohaplotypes (PS1–PS6) of *R. austrobrasilensis* across three chromosomes, revealing extensive rearrangements and inversions while retaining conserved sets of centromeric arrays; (c) across three phased haplotypes of *R. gaudichaudii*, illustrating largely conserved centromeric array positions with localised rearrangements between haplotypes; (d) inter-chromosomal *Tyba*-array synteny in *R. ridleyi*, highlighting redistribution of conserved centromeric arrays consistent with chromosome rearrangements and fusion events after whole genome duplication; (e) within *R. pubera* genome showing the large-scale conservation of *Tyba*-array synteny after the two rounds of WGD, as depicted by the schematic of ancestral synteny blocks below it; (f) within *R. tenerrima*, showing extensive reorganisation between duplicated chromosome sets of subgenome A and B, while preserving array identity. Each chromosome to scale.

**Extended Data Fig. 5:**
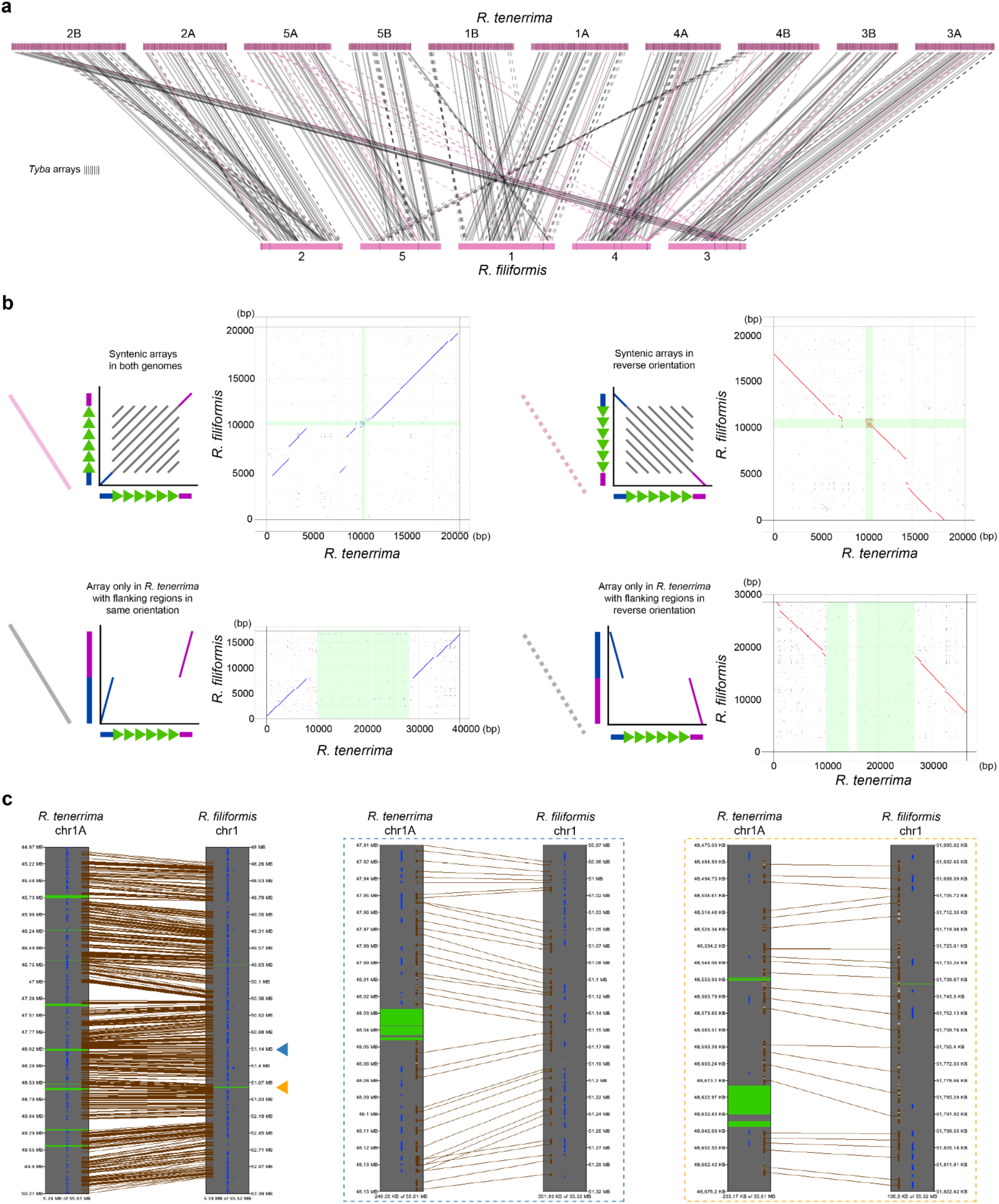
Synteny-guided reconstruction reveals recent loss of *Tyba* centromeric arrays in *R. filiformis*. (**a**) Chromosome-scale flanking *Tyba*-array synteny comparison between *R. tenerrima* and *R. filiformis*. Coloured bars represent homologous chromosomes, and connecting lines indicate conserved syntenic *Tyba* arrays. *Tyba* arrays are marked along chromosomes, highlighting their extensive presence in *R. tenerrima* and near-complete loss in *R. filiformis*. (**b**) Representative dot plots illustrating array matching outcomes used by the bidirectional tracking algorithm. Panels show syntenic *Tyba* arrays shared between both genomes, arrays conserved in reverse orientation, and arrays present only in *R. tenerrima* whose flanking regions are preserved in *R. filiformis* in either the same or reverse orientation. Green-shaded regions denote *Tyba* array annotations in one or both genomes. (**c**) Fine-scale SyMAP alignments between *R. tenerrima* and *R. filiformis* highlighting cases of complete (blue arrowhead and zoomed in blues dashed square) and incomplete (yellow arrowhead and zoomed in yellow dashed square) *Tyba* array erosion. Green blocks mark *Tyba* arrays present in *R. tenerrima* but largely absent in *R. filiformis*, while conserved flanking regions maintain collinearity (brown lines) and gene order (blue bars). Together, these analyses demonstrate a recent and extensive loss of *Tyba* centromeric arrays in *R. filiformis* that occurs without disruption of the surrounding genomic context, indicating that centromeric satellite erosion can proceed independently of large-scale chromosomal rearrangements.

**Extended Data Fig. 6:**
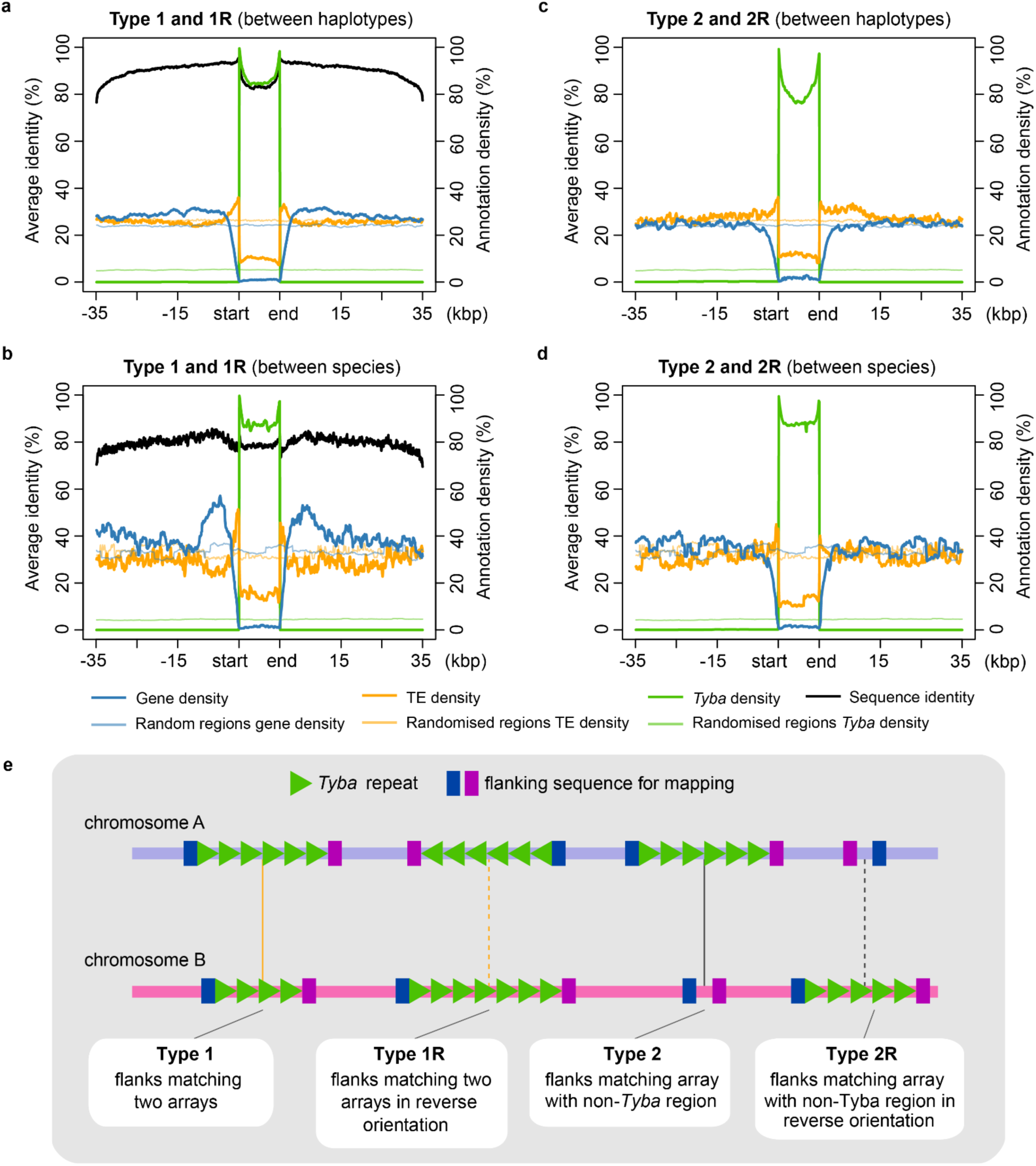
Sequence composition and conservation across syntenic *Tyba* array matches. (**a–b**) Average sequence identity and annotation density profiles centred on *Tyba* arrays for Type 1 and Type 1R matches, calculated between haplotypes (**a**) and between species (**b**). Between species, regions flanking retained array pairs (Type1/1R) show a significant enrichment of genes within 2–10 kbp of array boundaries compared with haplotype-level comparisons (47.50% versus 37.07%, Mann–Whitney U test *p* = 0.018), a difference not observed further from arrays (10–20 kbp: 38.44% versus 36.97%, *p* = 0.54). Sequence identity within *Tyba* arrays is significantly lower than in surrounding regions (81.04% within arrays versus 85.66% in the 15 kbp flanking Type 1/1R regions, Wilcoxon signed-rank test *p* = 0.002). (**c–d**) Equivalent profiles for Type 2 and Type 2R matches, where flanking regions of *Tyba* arrays in one genome align to syntenic non-*Tyba* regions in the other, calculated between haplotypes (**c**) and between species (**d**). In contrast to Type 1/1R loci, no significant gene enrichment is detected at Type 2/2R loci (2–10 kbp: 30.42% versus 32.17%, *p* = 0.689; 10–20 kbp: 38.44% versus 36.97%, *p* = 0.644). Across all match types, array boundaries are enriched for short, fragmented TEs, with significantly higher TE density in the first 2 kbp flanking arrays compared with the subsequent 2–10 kbp in Type 1/1R regions (37.78% versus 28.99%, Wilcoxon signed-rank test *p* < 0.001) and a smaller but significant difference in Type 2/2R regions (37.25% versus 35.22%, *p* = 0.01). Plots show average sequence identity (black), *Tyba* density (green), gene density (blue) and TE density (orange), with corresponding randomised controls indicated by lighter lines. (**e**) Schematic illustrating the four array-matching categories defined by flanking sequence similarity and orientation: Type 1, orthologous arrays with conserved flanks; Type 1R, orthologous arrays in reverse orientation; Type 2, arrays aligned to syntenic non-*Tyba* regions; and Type 2R, equivalent matches in reverse orientation.

**Extended Data Fig. 7:**
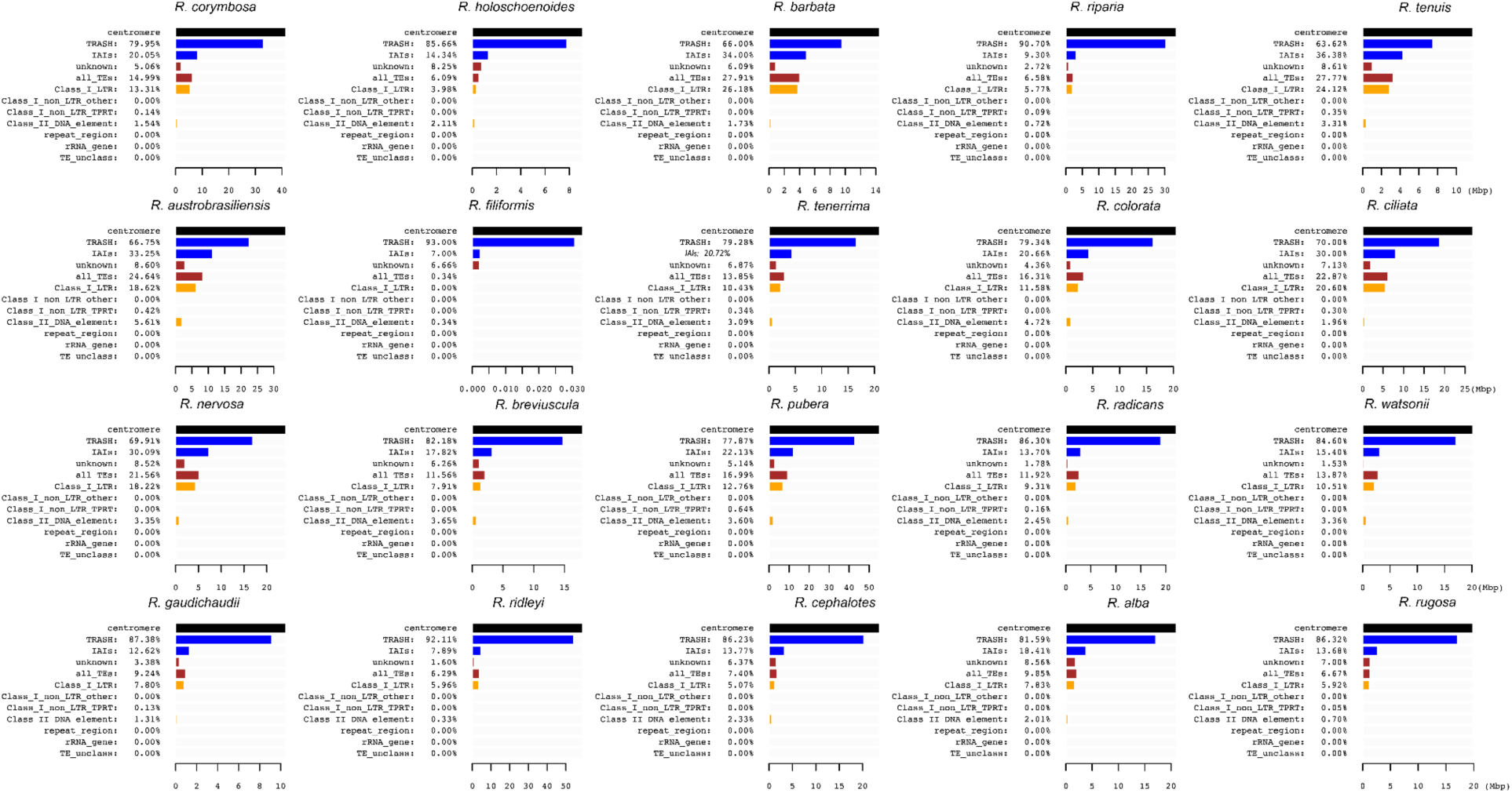
Composition of centromeric space across *Rhynchospora* species. Horizontal bar plots summarise the sequence composition of *Tyba*-defined centromeric regions for each of the 20 *Rhynchospora* species. Bars indicate the genomic contribution in Mbp of *Tyba* satellite DNA (TRASH) and intra array interruptions (IAIs), which are further split into transposons (all_TEs) and DNA of unknown origin (with the latter possibly including low quality Tyba and transposon sequences that have not been annotated by our pipelines). Transposon sequences are classified into Class_I_LTRs (i.e. different types of LTR retrotransposons), Class_I_nonLTR_TPRT (i.e. non-LTR retrotransposons like LINEs that undergo target-primed reverse transcription), Class_I_nonLTR_other (e.g. Penelope and Tyrosine Recombinase elements), Class_II_DNA_element (e.g. transposons like Mutators that have terminal inverted repeats, and also Helitrons), and transposon or other type of repetitive DNA (TE_unclass and repeat_region) that was not assigned to a specific type by EDTA. Absolute centromeric sequence length is shown on the *x*-axis for each species. Across the genus, centromeric regions are consistently dominated by *Tyba* repeats, with variable but generally limited contributions from transposons. Species-specific differences in centromeric composition and size are evident, including the markedly reduced centromeric space in *R. filiformis*.

**Extended Data Fig. 8:**
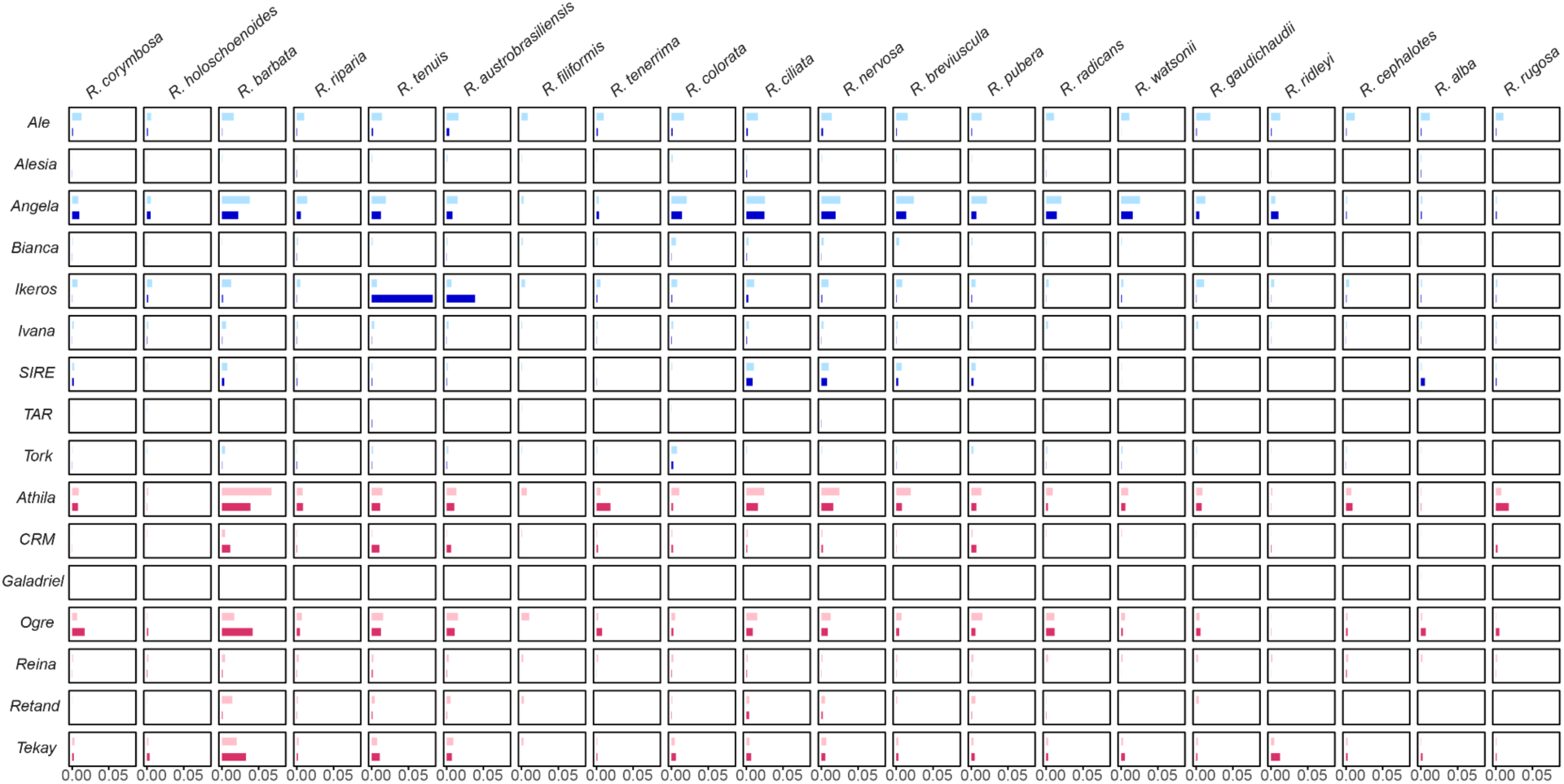
Diversity of LTR lineages inserted within *Tyba* centromeric arrays across *Rhynchospora*. Stacked bar plots summarise the composition and abundance of LTR retrotransposon lineages within *Tyba*-defined centromeric units (dark blue for Ty1/Copia lineages, dark red for Ty3/Gypsy lineages) and in the rest of the genome (light blue for Ty1/Copia lineages, light red for Ty3/Gypsy lineages) across 20 *Rhynchospora* species. The distribution reveals both species-specific centromeric enrichment of certain LTR lineages (for example, *Ogre* in *R. barbata* and *Ikeros* in *R. tenuis* and *R. austrobrasiliensis*), *Rhynchospora*-wide lack of centromere invasion (i.e. within the centromeres Ale represents ∼2.7% of the total fraction of all LTR lineages combined across all species versus ∼16.1% for outside the centromeres), and invasion dynamics for some lineages that broadly mirror their presence outside the centromeres (e.g. Angela and Athila show similar proportions of genome and TE space inside and outside the centromeres). Together, these patterns highlight the complex colonisation dynamics (both genus-consistent and species-specific) of the *Tyba*-based centromeric space of *Rhynchospora* species by different LTR retrotransposon lineages. See **Supplementary Dataset 6** for the proportion of every LTR lineage per species and for all genomes combined, both inside and outside the centromeres.

**Extended Data Fig. 9:**
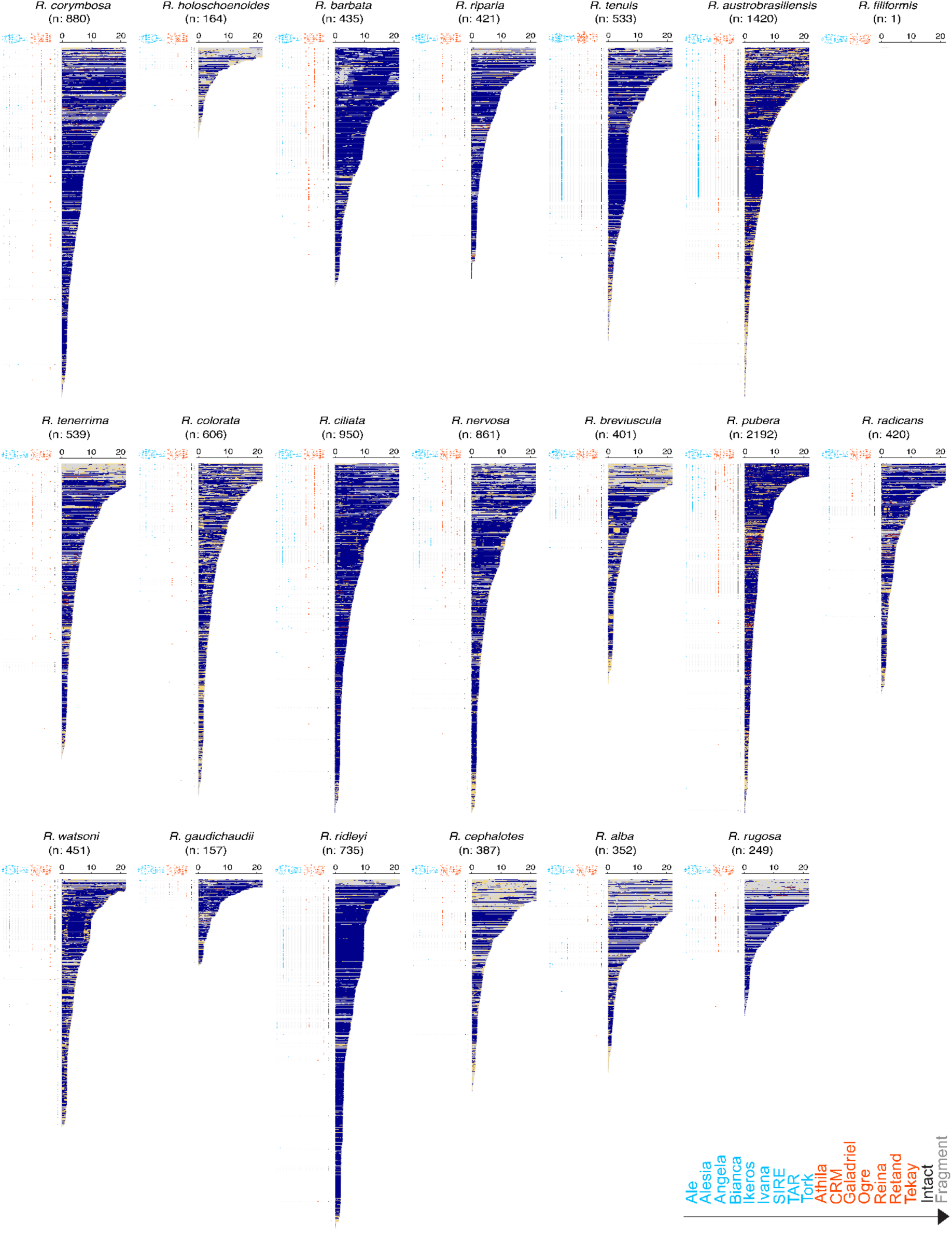
Composition and sequence characteristics of centromeric *Tyba* intra-array interruptions (IAIs) across *Rhynchospora* species. IAIs are ordered according to their physical length in kilobase pairs (kbp), and coloured according to their transposon content along their sequence. IAI lengths are capped at 20 kbp for visual clarity (maximum length of an IAI can be 50 kbp, but the vast majority are <20 kbp). The presence of different LTR lineages (Ty1/Copia: blue, Ty3/Gypsy: red) is included on the left of every IAI. Two additional columns indicate intact elements (black) and their fragments (dark grey) to emphasize the large number of IAIs that are composed entirely by a single intact element in a small number of species (Ikeros in *R. tenuis* and *R. austrobrasiliensis*, Angela and Tekay in *R. ridleyi*), which are consistent with recent transposon invasion.

**Extended Data Fig. 10:**
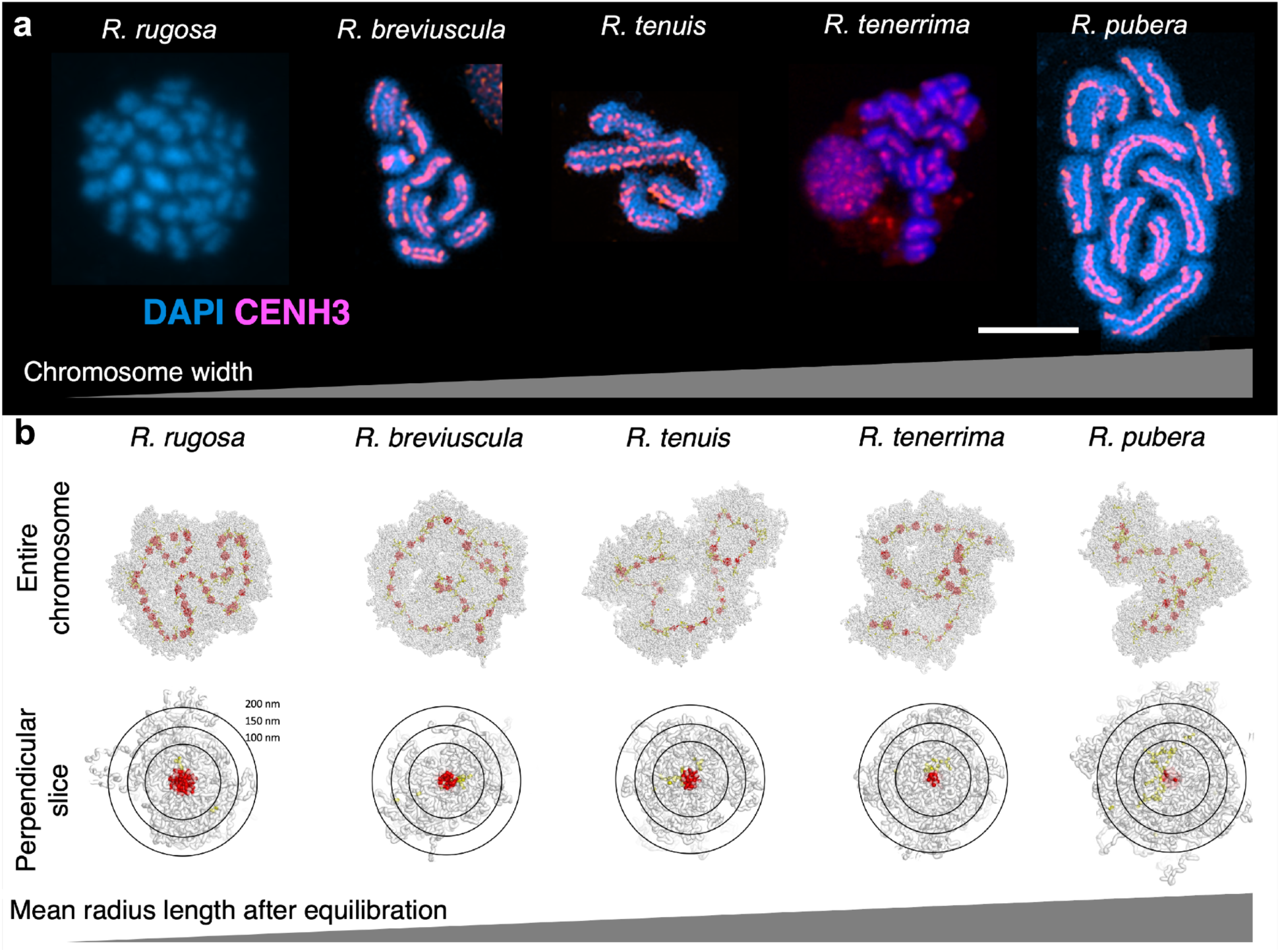
Holocentromere organisation and inter-array spacing scale chromatid thickness and loop architecture. **(a**) representative metaphase chromosomes from five *Rhynchospora* species stained with DAPI (blue) and CENH3 (magenta), illustrating progressive increases in chromosome width across species. Scale bar, 5 µm. (**b**) corresponding polymer simulations of holocentric chromosomes parameterised with species-specific *Tyba* array spacing. Upper panels show equilibrated chromosome models highlighting the distribution of centromeric units along the chromatid axis. Lower panels show perpendicular cross-sections of simulated chromatids, with concentric circles indicating distances from the chromatid centre (100, 150 and 200 nm). Increasing inter-array spacing is associated with larger chromatin loops and increased mean chromatid radius after equilibration. Together, the experimental and modelling data demonstrate that variation in *Tyba* inter-array spacing contributes directly to differences in chromatid thickness and higher-order chromosome architecture across *Rhynchospora*.

**Supplementary Fig. 1:**
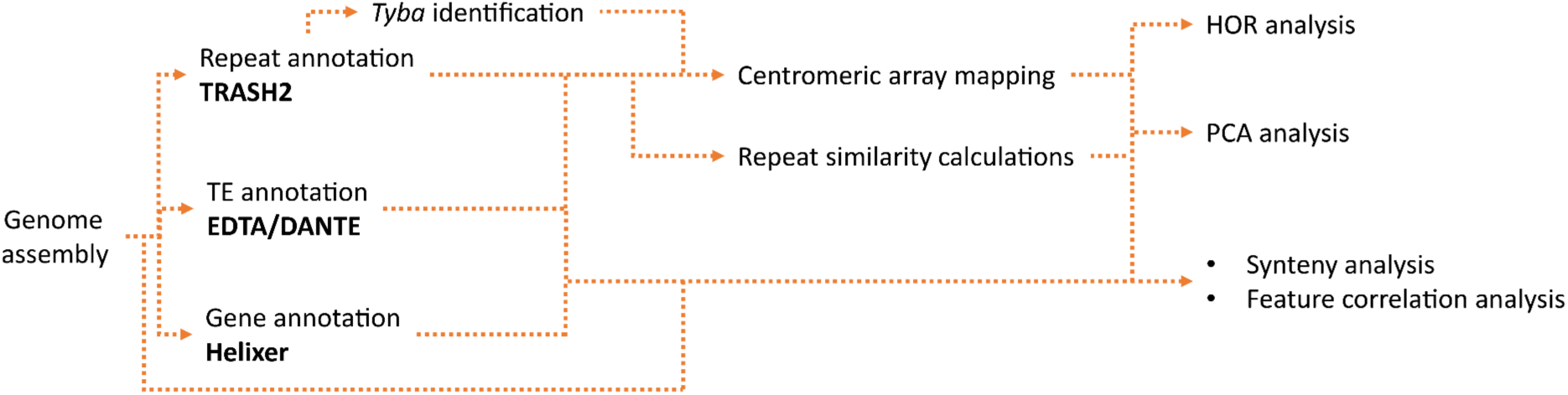
Overview of the analytical pipeline used to characterise repeat-based holocentromeres in *Rhynchospora*. Schematic outlining the computational workflow starting from chromosome-scale genome assemblies. Repeat annotation was performed using TRASH2 to identify *Tyba* satellite repeats and EDTA/DANTE for transposable element annotation, while gene models were generated using Helixer. These annotations were integrated to map centromeric *Tyba* arrays, calculate repeat similarity, and perform higher-order repeat (HOR) and principal component analyses. Combined repeat, gene and transposon annotations were further used for synteny and feature correlation analyses across haplotypes and species.

**Supplementary Fig. 2:**
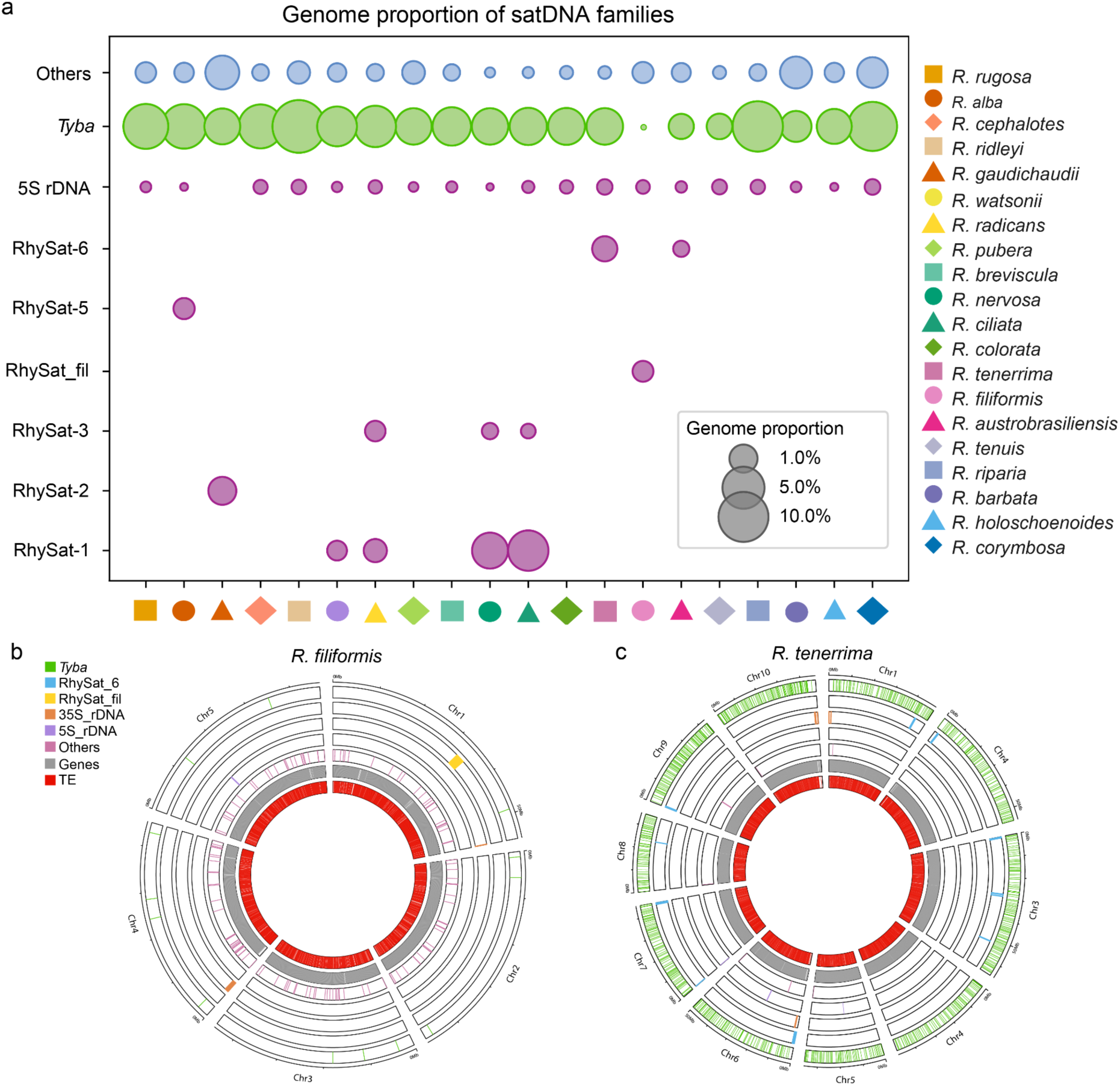
Satellite DNA evolution in *Rhynchospora*. (**a**) Genome proportion of different families of satDNA among the 20 *Rhynchospora* species. *Tyba* (green) is the only satellite to found across all Rhynchospora species, while other abundant families (purple), mostly lineage-specific, are shown separately (RhySat-#), while the rest of less abundant families of satDNA are plotted together for each species as “Others” (blue). **(b–c)** Genome-wide distribution of the main families of repetitive elements (satDNA and TEs), and genes in (**b**) *R. filiformis,* and (**c**) *R. tenerrima*.

**Supplementary Fig. 3:**
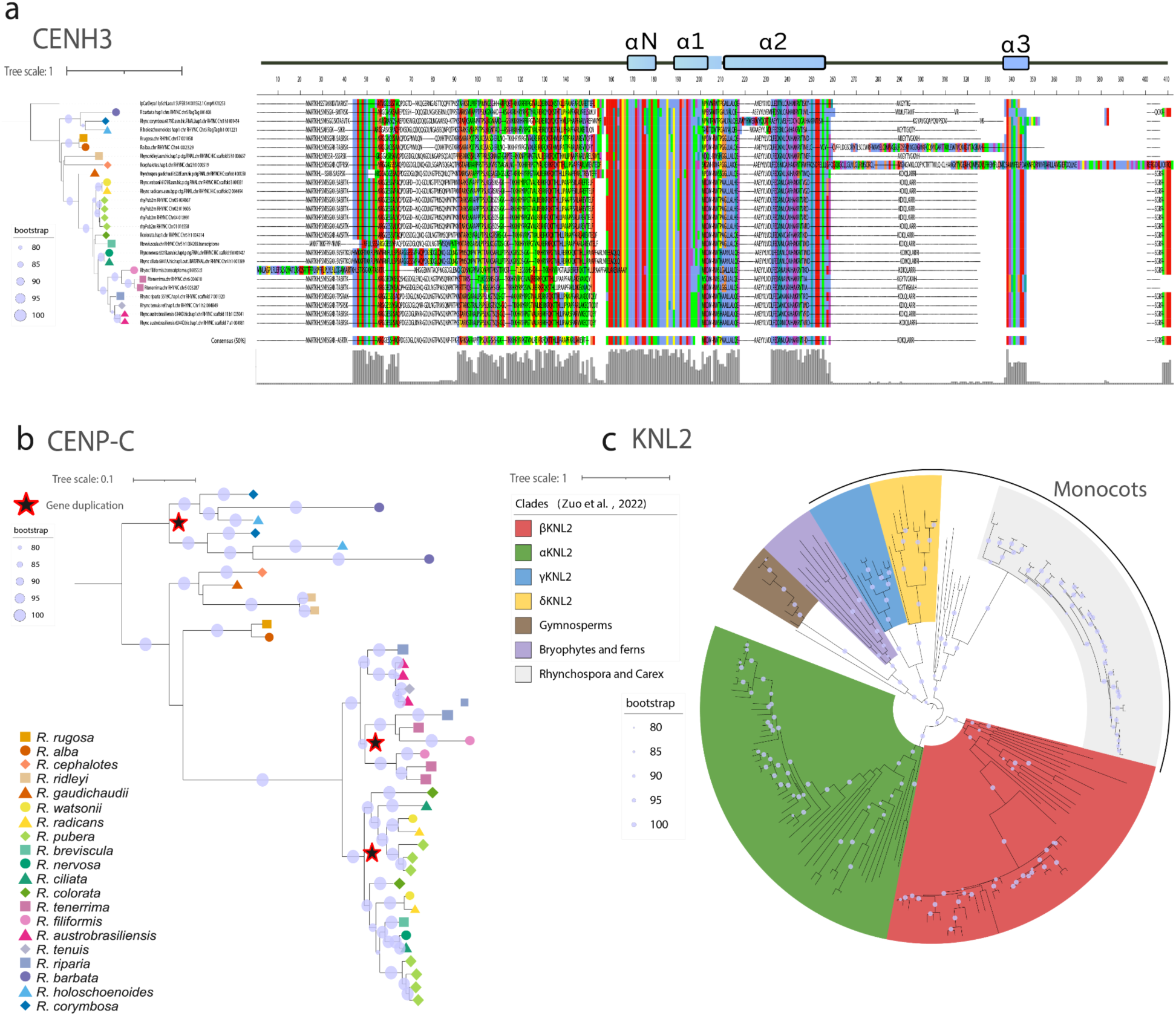
Characterisation of CENH3, CENP-C and KNL2 across 20 *Rhynchospora* species. (**a**) CENH3 protein alignment and maximum likelihood phylogenetic tree. Conserved histone H3 alpha helices (α1, α2, α3) are highlighted. (**b**) CENP-C phylogenetic tree showing gene duplication events (red stars). (**c**) Rhynchospora KNL2 and its phylogenetic relationships across plant lineage, with sequences retrieved from *Zuo et al*. Major clades are color-coded: monocots (yellow), KNL2 variants (βKNL2, αKNL2, γKNL2, δKNL2), gymnosperms (green), bryophytes and ferns (gray), and Rhynchospora/Carex (brown/red). Bootstrap support values (80-100%) are indicated by circle size across all trees.

**Supplementary Fig. 4:**
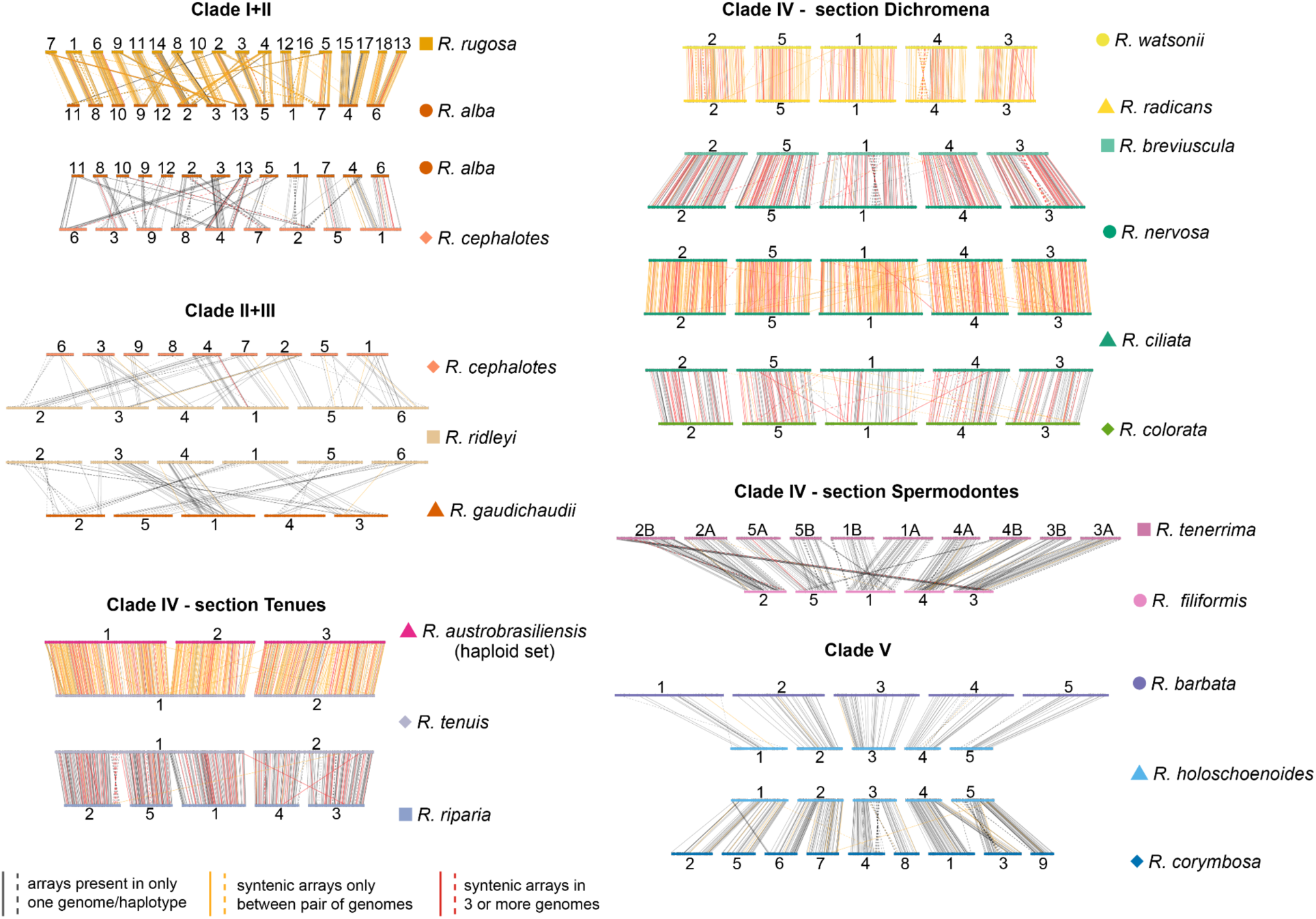
Lineage-specific patterns of *Tyba* array synteny across the *Rhynchospora* phylogeny. Synteny-guided comparisons of *Tyba* centromeric arrays across major *Rhynchospora* clades, organised according to the phylogenetic classification of Costa et al. 2025. Panels show representative species comparisons within Clades I+II, II+III, IV (sections *Dichromena*, *Tenues* and *Spermodontes*), and Clade V. Horizontal bars represent chromosomes ordered by synteny groups, and connecting lines indicate orthologous *Tyba* arrays identified by flanking sequence conservation. Line colours denote array-sharing categories: arrays present in only one genome or haplotype (grey), syntenic arrays shared between a pair of genomes (orange), and arrays conserved across three or more genomes (red). Across clades, patterns range from extensive conservation of array positions within closely related lineages to pronounced lineage-specific gains, losses and rearrangements, illustrating how *Tyba* array turnover scales with phylogenetic divergence while preserving chromosome-wide centromere organisation.

**Supplementary Fig. 5:**
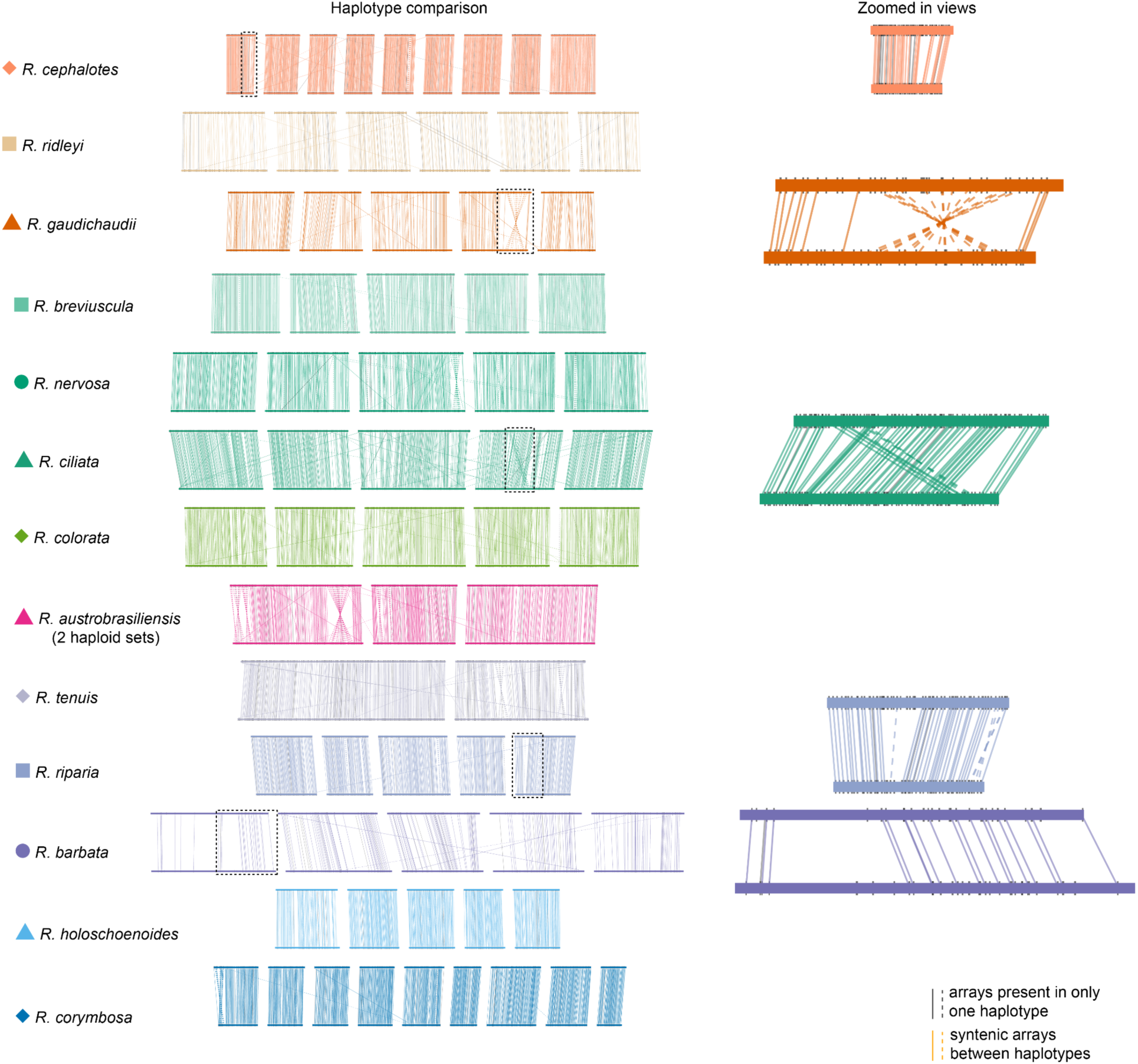
Conservation and local rearrangement of *Tyba* centromeric arrays across haplotypes in *Rhynchospora*. Haplotype-level comparisons of *Tyba* array positions across multiple *Rhynchospora* species. For each species, horizontal bars represent chromosomes from different haplotypes, and connecting lines indicate syntenic *Tyba* arrays identified by conserved flanking sequences. Dashed boxes highlight regions shown at higher resolution in the zoomed-in panels (right), which illustrate examples of local rearrangements, including inversions and spacing variation, while retaining overall array collinearity. Across species, *Tyba* arrays are largely conserved in position within haplotypes, with divergence mainly reflecting local structural changes rather than wholesale relocation or loss of centromeric units.

**Supplementary Fig. 6:**
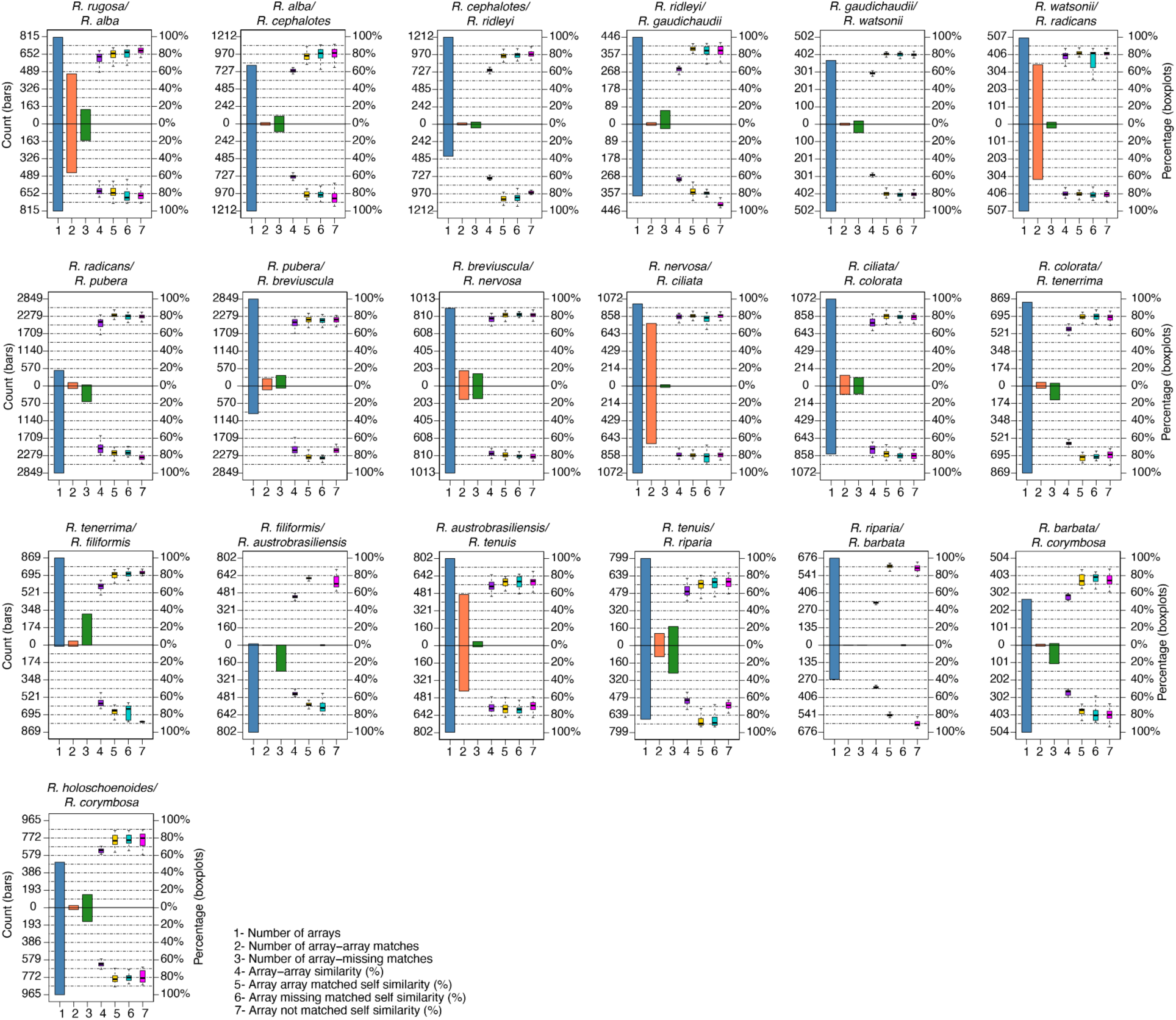
Quantitative summary of *Tyba* array conservation and turnover across pairwise species comparisons. Mirrored bar and boxplot representations summarise pairwise comparisons of *Tyba* arrays between species. For each species pair, bars (left axis) show the total number of arrays (1), the number of array–array matches (2), and the number of array–missing matches (3), reflecting array conservation and loss. Boxplots (right axis) show percentage similarity metrics, including array–array similarity (4), self-similarity of matched arrays (5), self-similarity of arrays with missing matches (6), and self-similarity of unmatched arrays (7). Together, these comparisons quantify the extent of *Tyba* array retention, divergence and loss across species, highlighting pronounced interspecific variability in centromeric array conservation despite shared genomic synteny.

**Supplementary Fig. 7:**
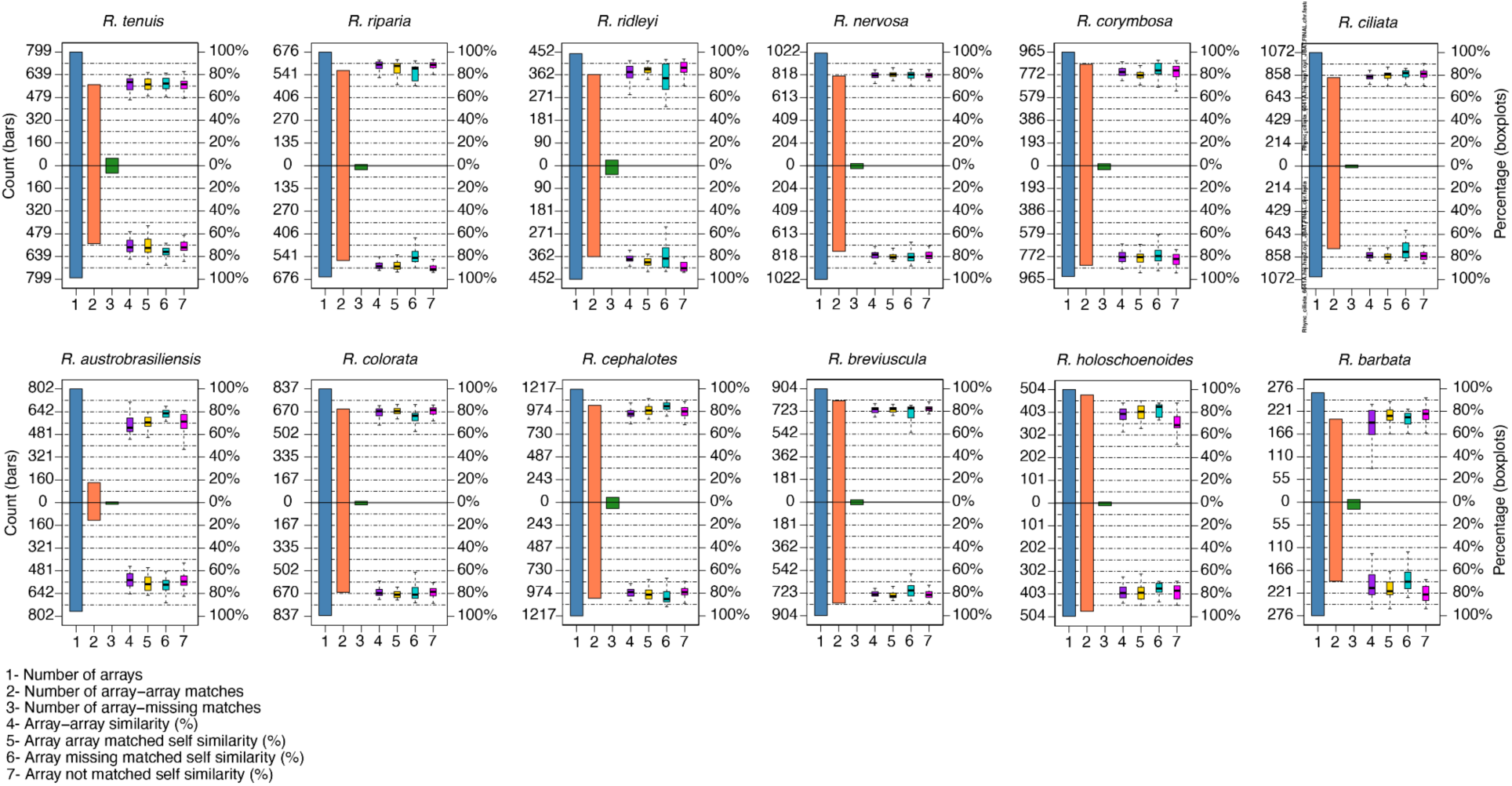
Quantitative summary of *Tyba* array conservation and turnover across haplotype comparisons. Mirrored bar and boxplot representations summarise haplotype-level comparisons of *Tyba* arrays within individual *Rhynchospora* species. For each species, bars (left axis) indicate the total number of arrays (1), the number of array–array matches between haplotypes (2), and the number of array–missing matches (3). Boxplots (right axis) show percentage similarity metrics, including array–array similarity (4), self-similarity of matched arrays (5), self-similarity of arrays with missing matches (6), and self-similarity of unmatched arrays (7). These analyses reveal high positional conservation of centromeric arrays within species, accompanied by moderate variation in array copy number and local organisation between haplotypes.

**Supplementary Figure 8:**
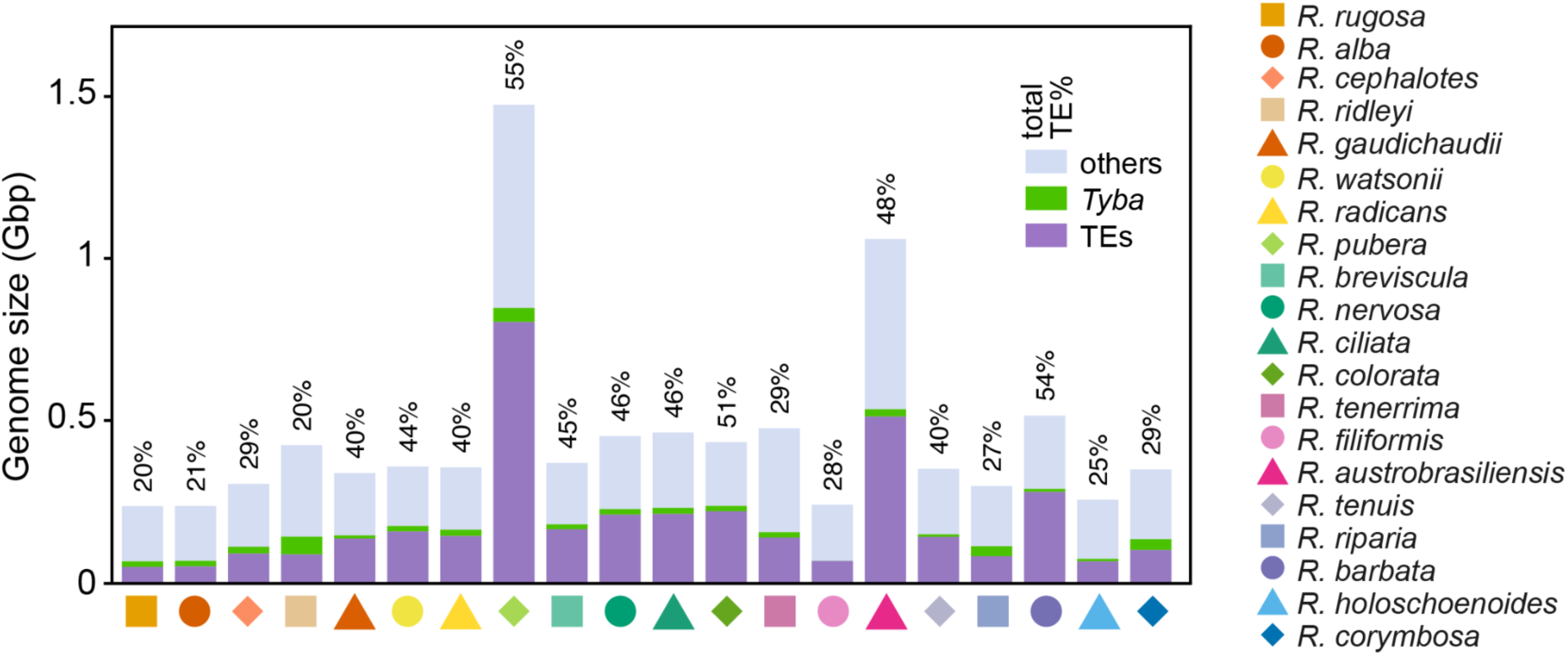
Correlation between total repeat abundance and genome size across the *Rhynchospora* genus. Stacked bar plots showing total genome size for each *Rhynchospora* species, partitioned into transposable elements (TEs; purple), *Tyba* satellite repeats (green), and other genomic components (light blue). Species are ordered phylogenetically and indicated by coloured symbols below the bars. Percentages above bars denote the proportion of the genome occupied by transposable elements. The figure highlights substantial variation in genome size across the genus and illustrates that genome expansion is primarily driven by TE content, while *Tyba* repeats contribute a relatively small but consistent fraction across species.

**Supplementary Fig. 9:**
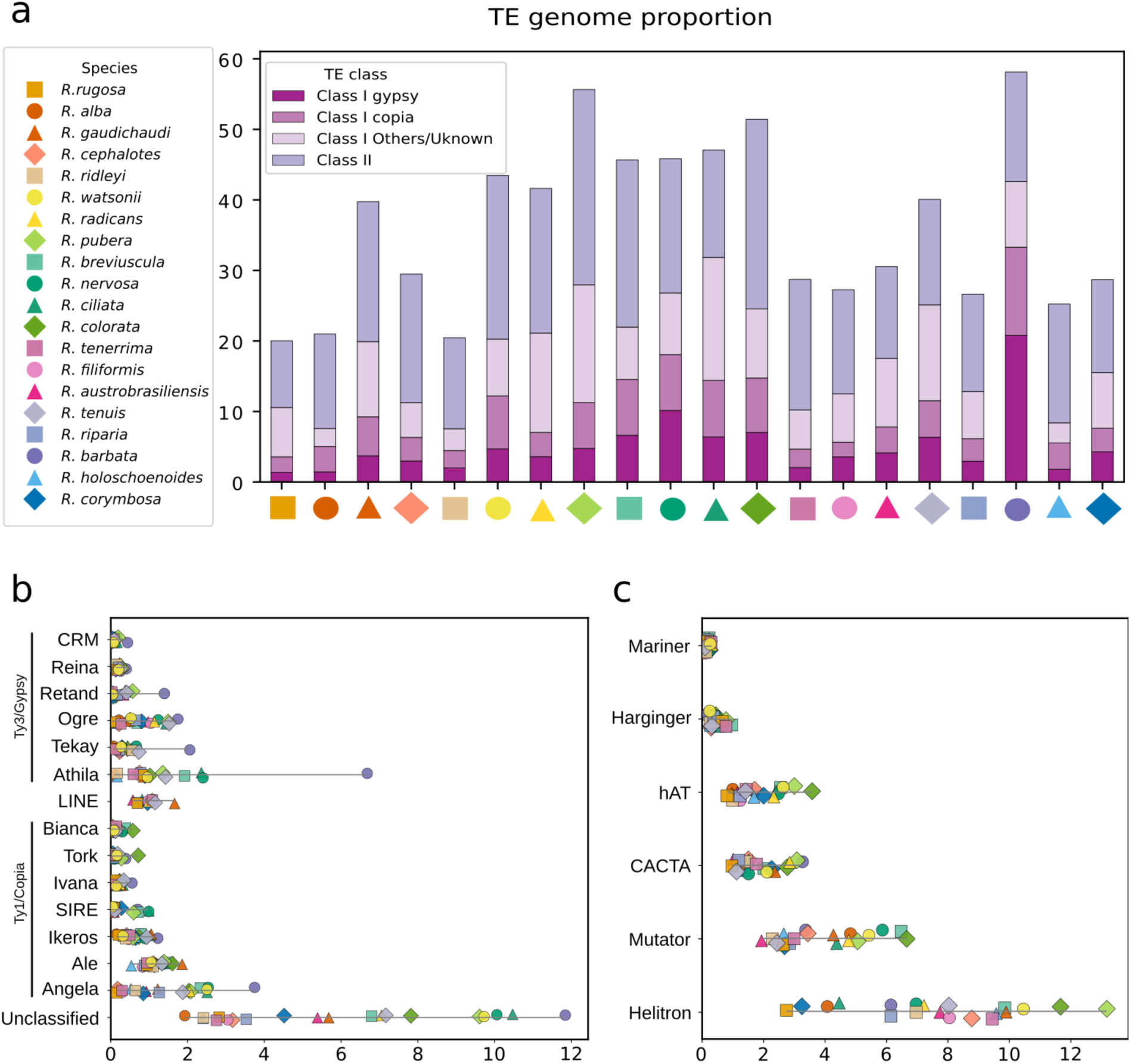
Genome-wide transposable element composition across 20 *Rhynchospora* species. (**a**) Proportion of Class I (retrotransposon) and Class II (DNA transposon) elements in each genome. Class I LTR retrotransposons predominate in most species, typically contributing more than half of the total TE fraction (∼50.1% for all species combined, see **Supplementary Dataset 5**). (**b**) Abundance of individual Class I LTR retrotransposon lineages. The Ty1/Copia lineage Angela and the Ty3/Gypsy lineage Athila are the most abundant (∼1.6% and ∼1.5% respectively for all genomes combined, see **Supplementary Dataset 6**), even though there are distinct species-specific differences both across the whole genome and also within the centromeric arrays only. (**c**) Abundance of Class II TE families, showing a high abundance of Helitron and broadly similar levels of hAT, CACTA, Mutator, Mariner and Harbinger across species. Together, these data indicate that a consistent set of TE lineages is shared across the genus, despite differences in total TE load.

**Supplementary Fig. 10:**
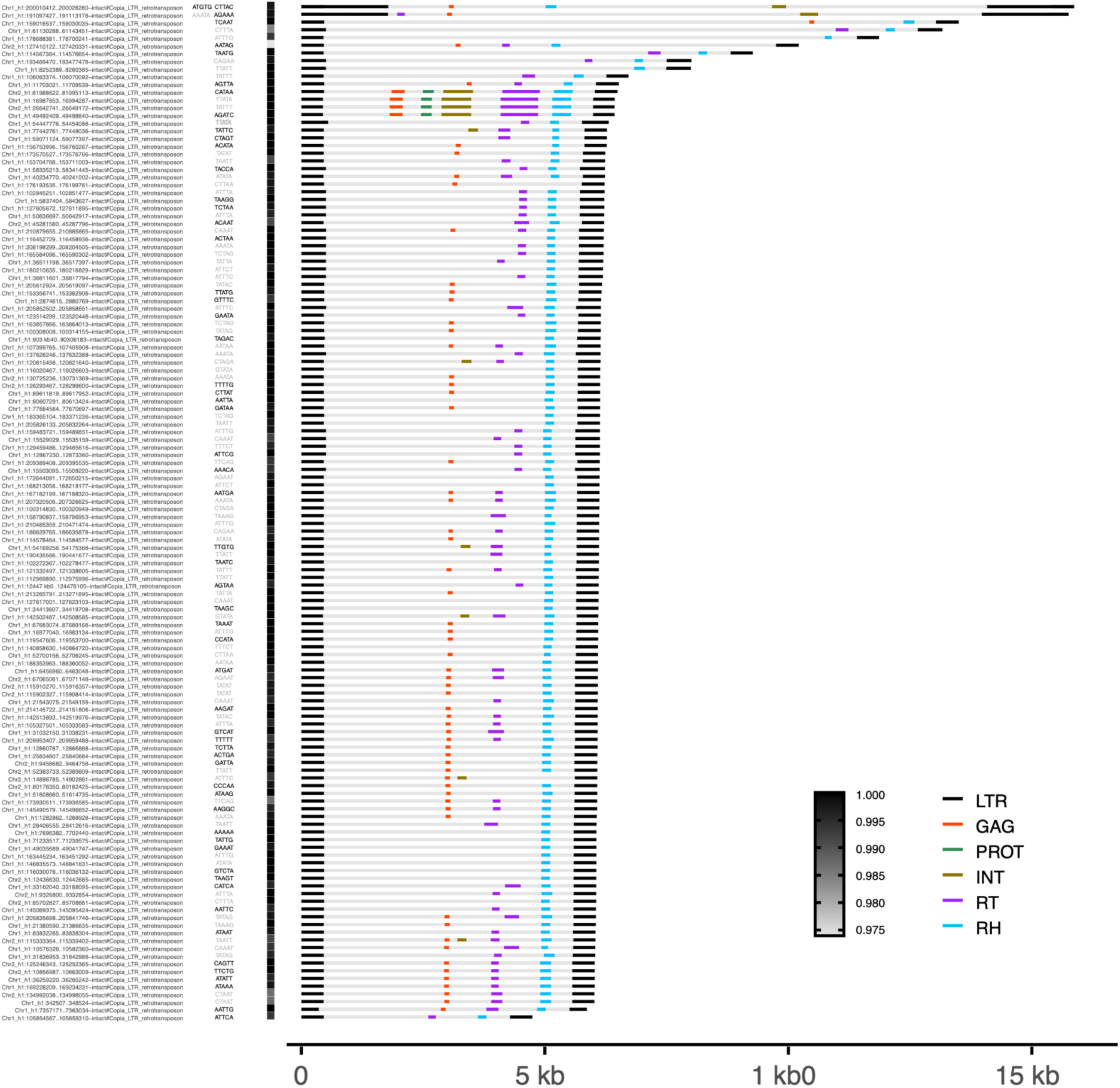
Structural validation of intact centrophilic Ikeros LTR retrotransposons inserted into *Tyba* centromeric arrays in *R. tenuis*. Each row represents an intact Ikeros element, scaled to its full length (*x*-axis). Black bars mark the long terminal repeats (LTRs), and coloured segments denote the core domains of the five canonical LTR retrotransposon genes (GAG; protease (PROT); integrase (INT); reverse transcriptase (RT); and RNaseH (RH). Greyscale shading indicates pairwise LTR sequence identity, confirming structural integrity of the elements. Nucleotide pentamers shown adjacent to each element represent target site duplications (TSDs): unique TSDs are shown in black, whereas shared TSDs occurring in more than one intact element are shown in grey, consistent with either chance recurrence or secondary duplication through non-TE recombination mechanisms. The consistent presence, organisation and identity of LTRs and coding domains demonstrate that Ikeros elements inserted into *Tyba* arrays are *bona fide* intact retrotransposons rather than annotation artefacts. We believe this is the first report for centrophilic adaptation of the Ikeros lineage in plants.

**Supplementary Fig. 11:**
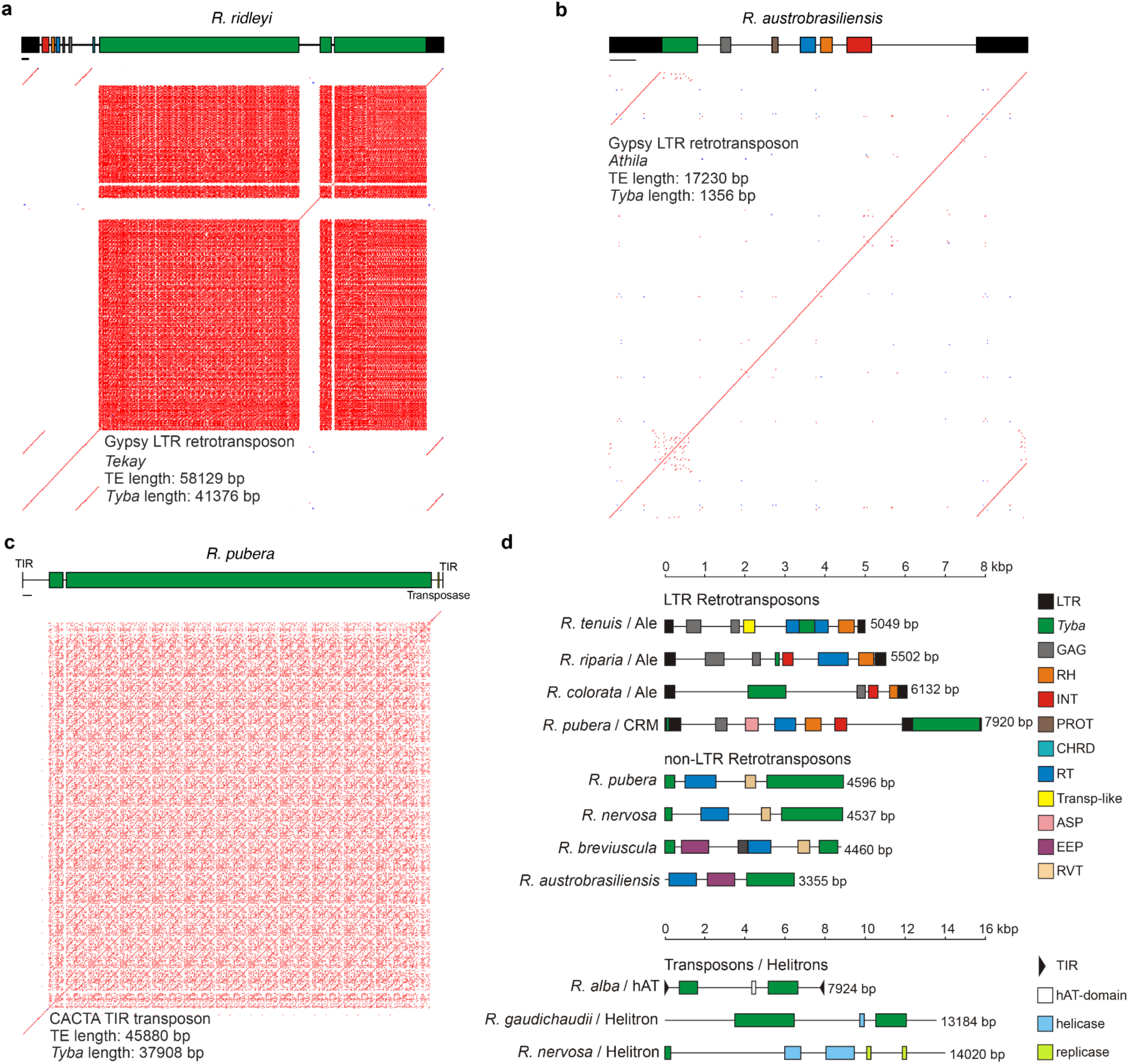
The association of *Tyba* repeats with TEs. (**a**–**c**) Representative sequence identity dot plots showing examples of *Tyba* insertions into intact TE sequences of different classes, showing variable degrees of structural preservation and integration. Each schematic illustrates *Tyba* arrays (green) embedded within a canonical LTR Ty3/Gypsy Tekay in *R. ridleyi* (**a**), a Ty3/Gypsy Athila in *R. austrobrasiliensis* (**b**) and a *CACTA TIR DNA transposon* in *R. pubera*, are shown as examples. Horizontal bars in **a–c** below the TE annotations correspond to 1 kbp. (**d**) Diversity of non-intact TE superfamilies carrying internal *Tyba* arrays, including LTR Ty1/Copia, Ty3/Gypsy, LINE, DNA transposon and Helitron elements, highlighting the broad spectrum of host–*Tyba* repeat interactions.

**Supplementary Fig. 12:**
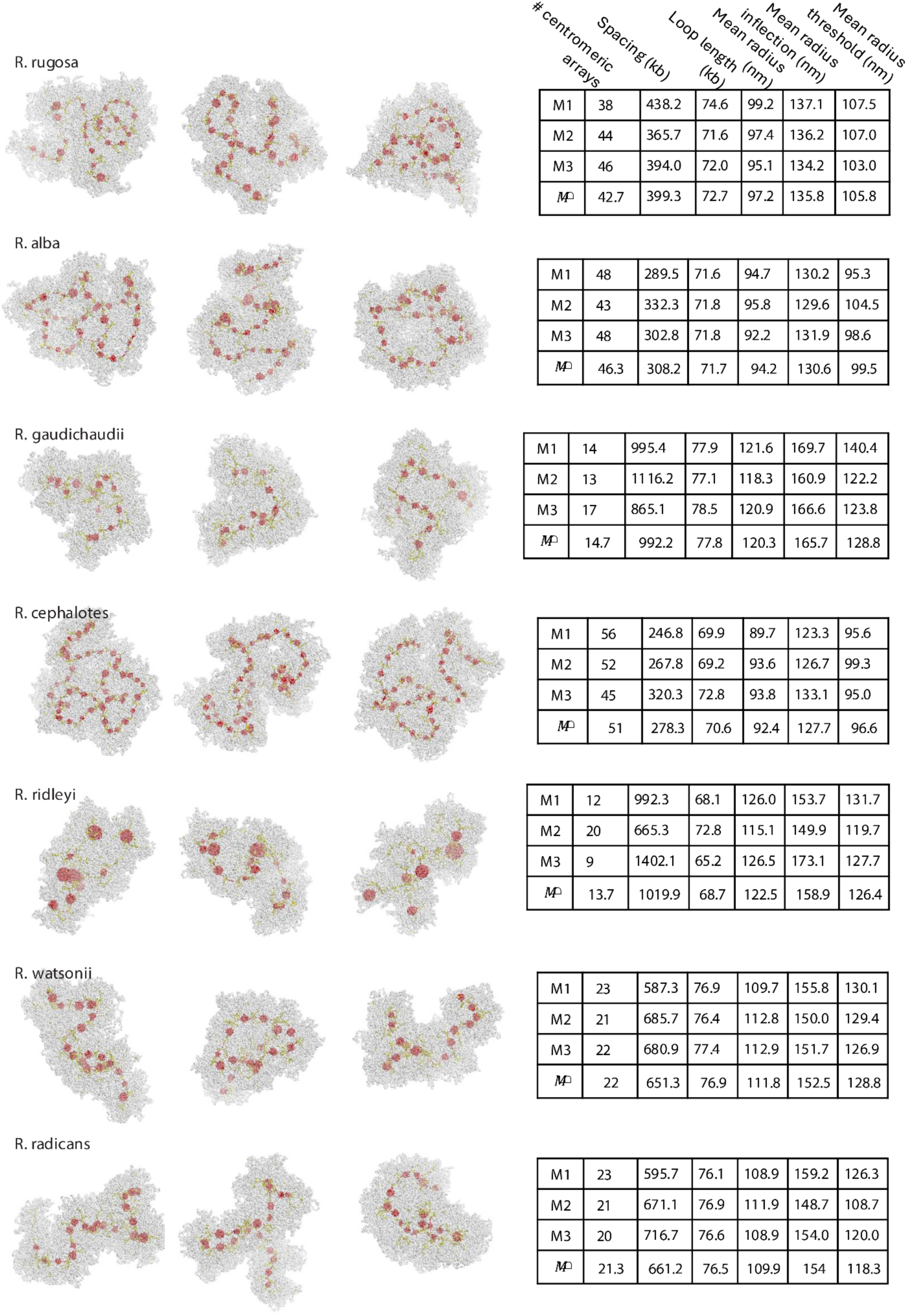
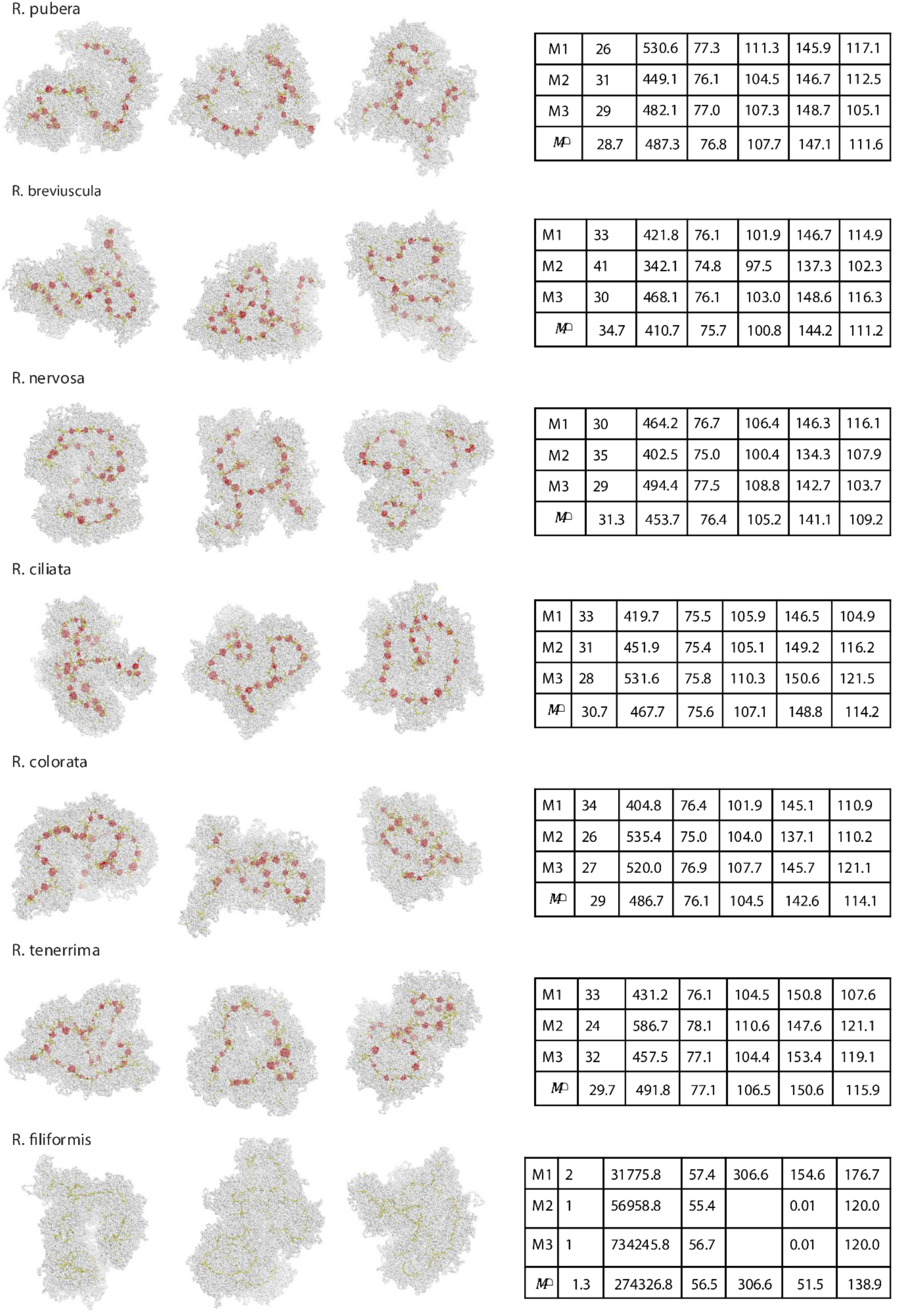
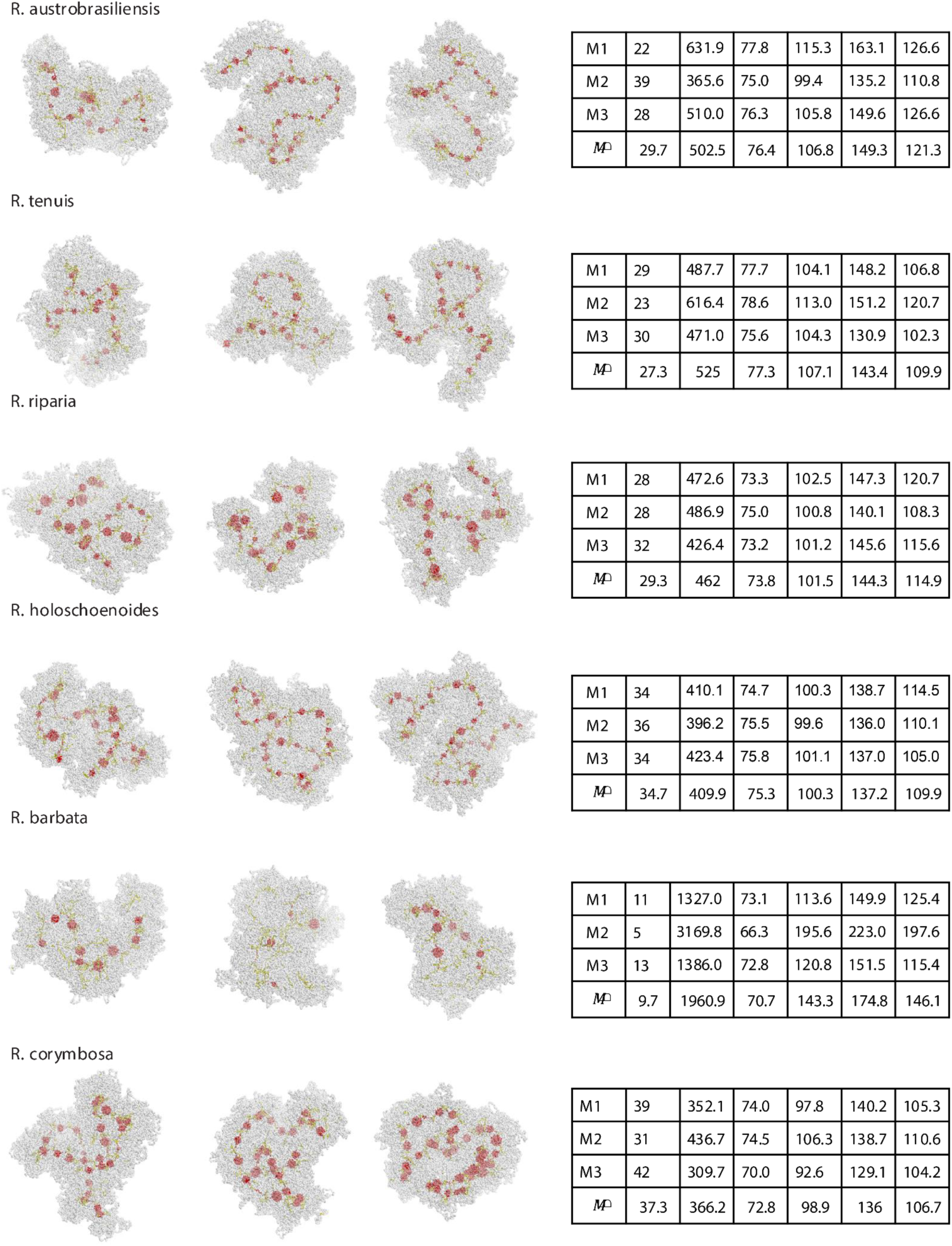
Simulated compaction of three different regions from the genomes of all 20 ***Rhynchospora* species.** All regions sum up 15 Mb. The last conformation of each is shown on the left, chromatin fiber in gray, centromeric nucleosomes in red and nucleosomes bound by loop extruders in yellow. Models 1 to 3 are shown from left to right. The table shows number of centromeric arrays, average spacing length, average loop length, mean radius calculated from the center view, and radius calculated as the inflection or as a threshold of a sigmoid calculated from top view, from left to right. Values for models 1 to 3 are shown from top to bottom, and the last line displays values averaged over the three models.

**Supplementary Table 1.**
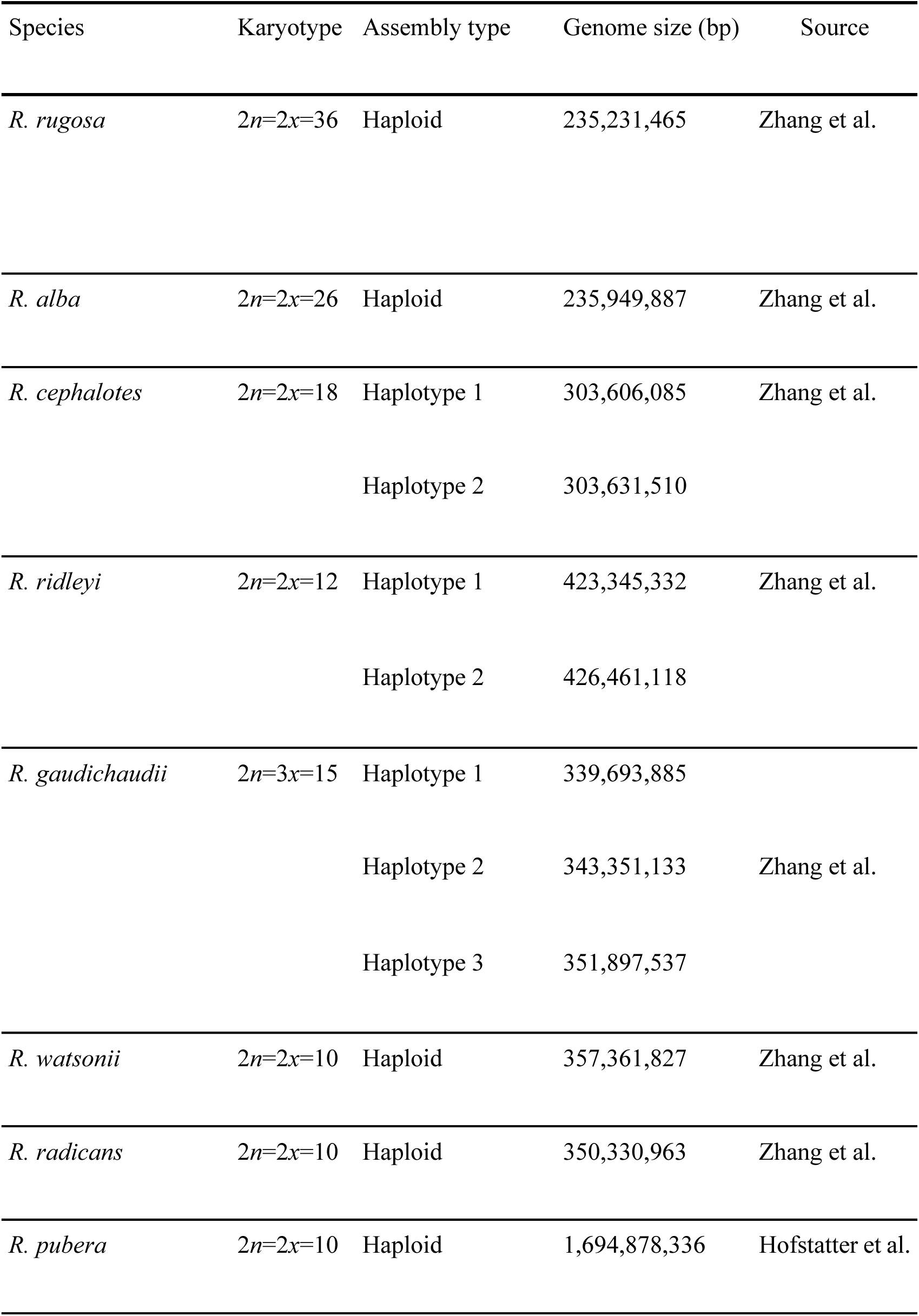
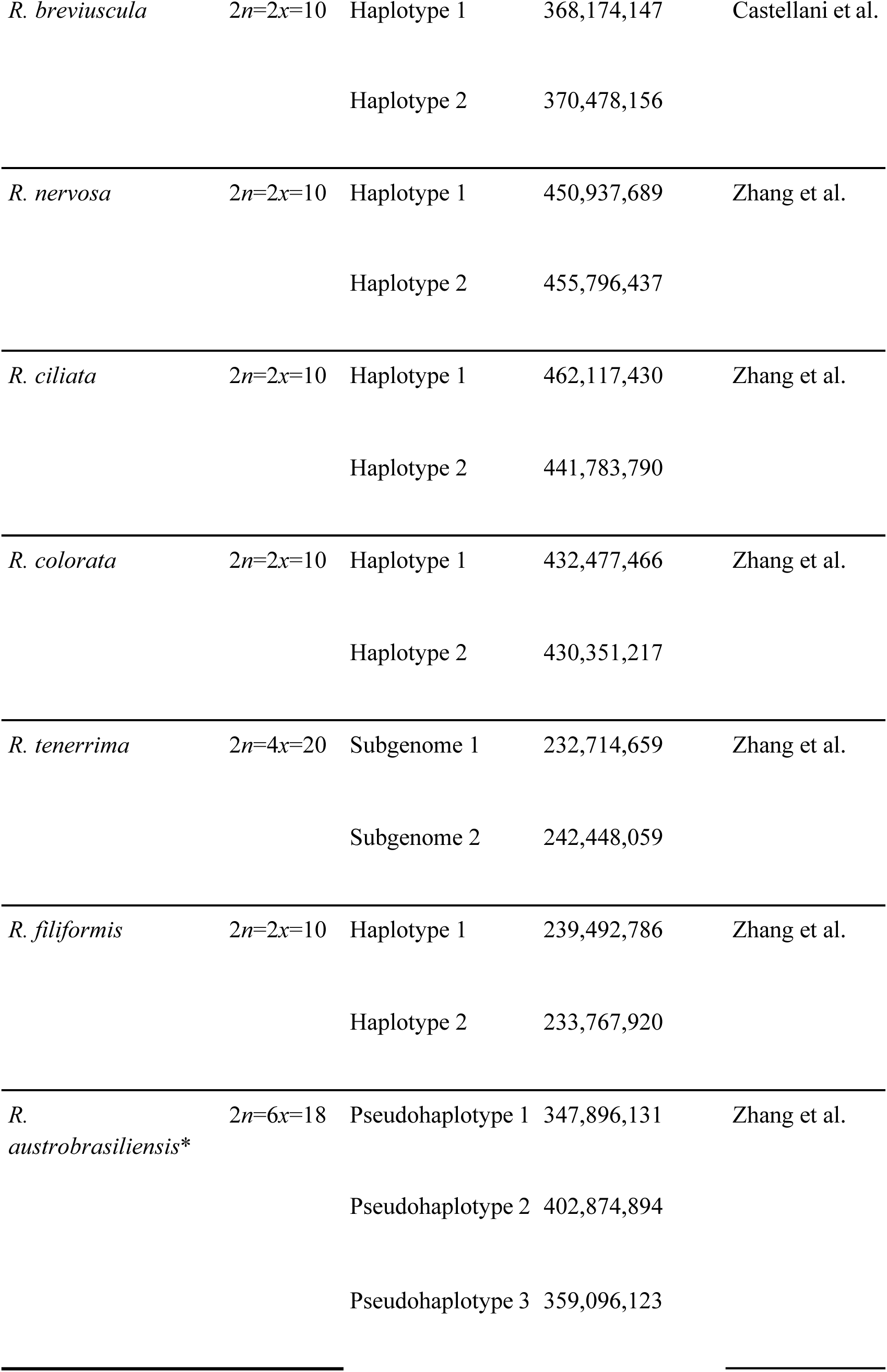
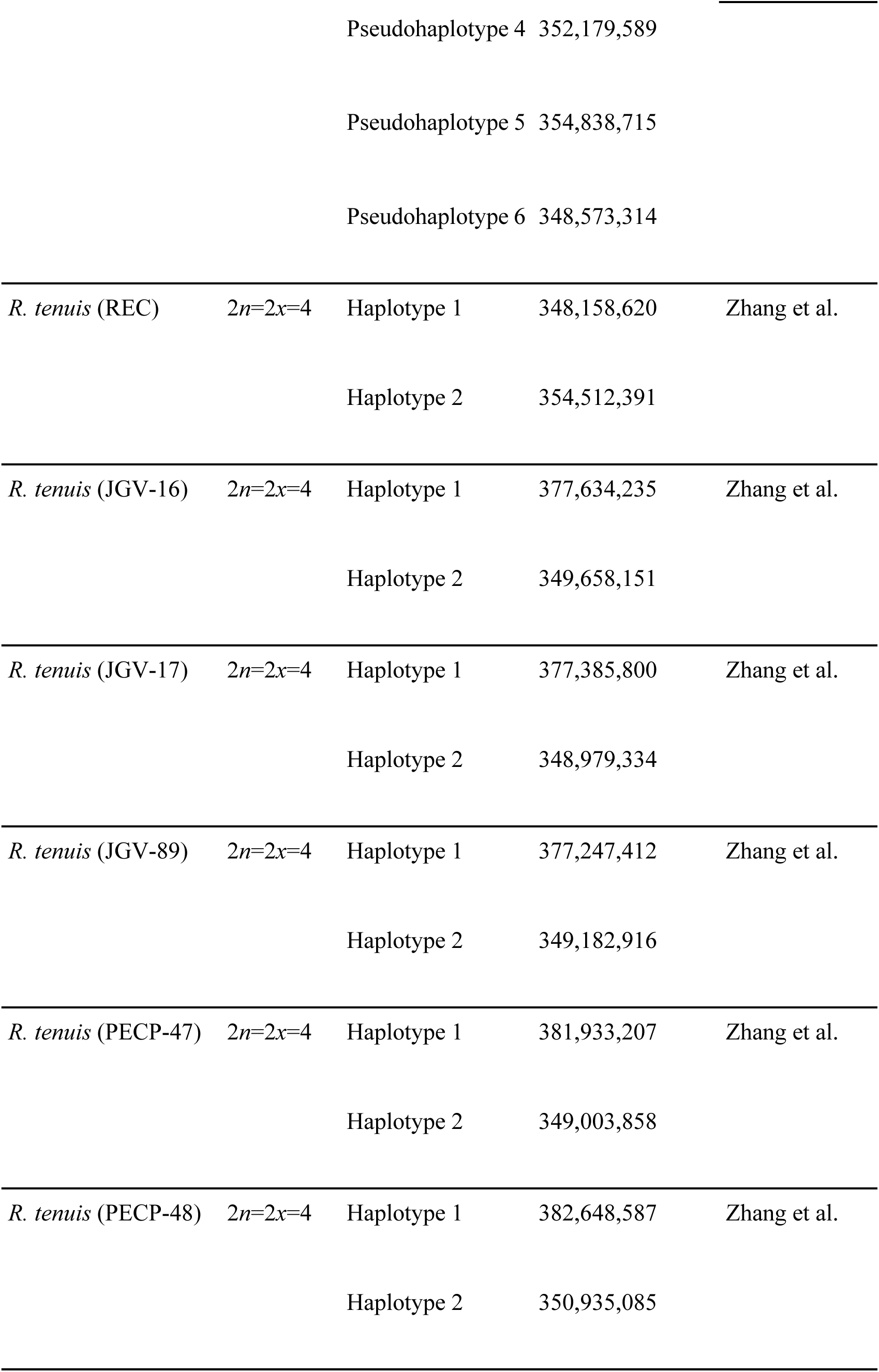
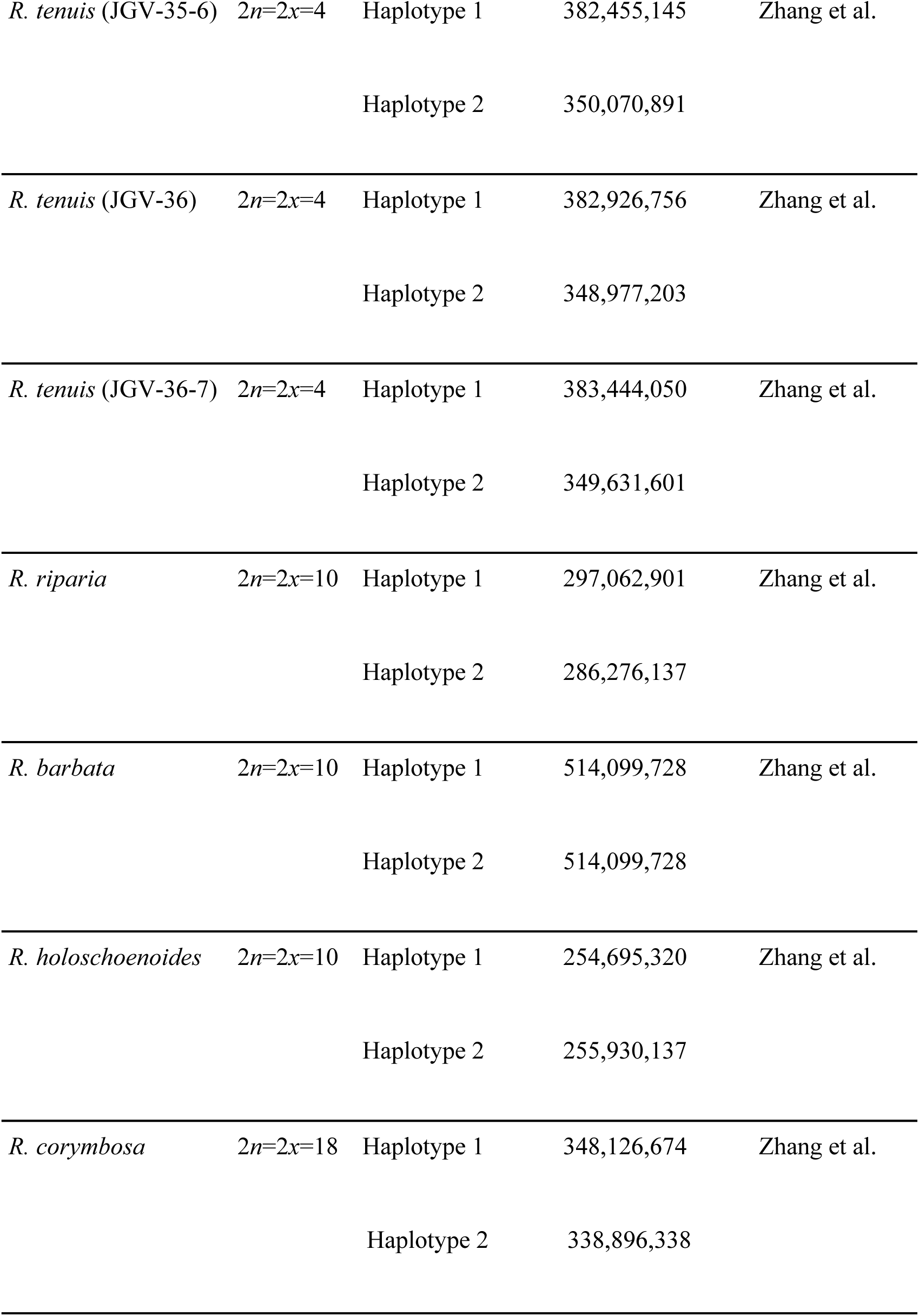
20 *Rhynchospora* species chromosome number, assembly type, genome size and assembly source.

## Supplementary Datasets

**Supplementary Dataset 1.** Metadata and annotation summary for repetitive elements across the *Rhynchospora* pangenome, including satellite repeats and transposable element classifications.

**Supplementary Dataset 2.** *Tyba* monomer annotations.

**Supplementary Dataset 3.** Synteny-aware *Tyba* array matching results between species pairs.

**Supplementary Dataset 4.** Synteny-aware *Tyba* array matching results between species haplotypes.

**Supplementary Dataset 5.** Genome-wide transposable element composition for all *Rhynchospora* assemblies.

**Supplementary Dataset 6.** Lineage-level annotation of LTR retrotransposons inside and outside *Tyba*-defined centromeric regions.

**Supplementary Dataset 7.** Extended results and parameters used for polymer simulations of holocentric chromosome architecture.

## Supplementary Movies

**Supplementary Movie 1.** Polymer simulation of a holocentric chromosome segment with short inter-array spacing based on *R. rugosa*, showing loop extrusion–mediated compaction that brings *Tyba* centromeric arrays into a linear holocentromeric axis.

**Supplementary Movie 2.** Polymer simulation of a holocentric chromosome segment with wide inter-array spacing based on *R. gaudichaudii*, illustrating reduced anchoring frequency during loop extrusion, formation of larger chromatin loops, and increased chromatid thickness.

## Notes

### Competing Interest Statement

The authors have declared no competing interest.

